# Neofunctionalised plastid translocon regulates chloroplast ageing in *Arabidopsis*

**DOI:** 10.64898/2026.09.29.755046

**Authors:** Pushan Bag, Panagiotis Lymperopoulos, Anne Lau, Bruno Peixoto, Charles Grundy, Sybille Kubis-Waller, Ramesh Patel, R. Paul Jarvis

## Abstract

Protein import across the chloroplast outer membrane is mediated by the TOC complex and regulated by GTP hydrolysis at two receptor GTPases, large and small. The former are the β-barrel-containing Toc159 family of proteins (Sun and Jarvis 2023; H. Li and Chiu 2010; Shi and Theg 2013a; Kessler and Schnell 2009). The smallest member of this family in higher plants is Toc90, which has long been regarded as functionally unimportant due to the absence of obvious phenotypes in early mutant studies (Kubis et al. 2004; Infanger et al. 2011a). It was hypothesised that Toc90 is a relic of an ancestral Toc159 protein from early green lineage organisms (Hofmann and Theg 2003; Reumann et al. 2005). Here, we challenge this view by combining genetic, biochemical, phylogenetic, and *in silico* structural analyses to show that Toc90 is not an ancient remnant, but rather a neofunctionalised derivative of Toc159, and likely the most recently evolved member of the family. We show that Toc90 physically associates with Toc33 as a part of a photosynthetic TOC complex. Furthermore, we identify Toc90 as a regulator of chloroplast ageing. Our findings redefine Toc90 as an active and specialised component of the chloroplast protein import machinery, playing a pivotal role in senescence, especially in reducing photosynthetic protein abundance. Together, these results highlight Toc90 as a promising target for manipulating chloroplast longevity and plant senescence, with potential to improve crop productivity and yield.

## Introduction

During eukaryotic evolution, the endosymbiotic organelles, plastids and mitochondria, transferred most of their ancestral genomes to the nucleus. Consequently, these organelles rely predominantly on imported nucleus-encoded, cytosol-synthesised precursor proteins for essential metabolic processes such as photosynthesis and respiration (Timmis et al. 2004). Such import is mediated by specialised translocon complexes spanning the organelles’ double bounding membranes (Sun and Jarvis 2023; H. Li and Chiu 2010; Shi and Theg 2013a; Kessler and Schnell 2009; Chen et al. 2000). In plastids, protein translocation occurs through two sequentially acting systems: the TOC (Translocon at the Outer Chloroplast membrane) and TIC (Translocon at the Inner Chloroplast membrane) complexes (Chotewutmontri et al. 2017; Sun and Jarvis 2023).

In land plants, the plastid protein import machinery has diversified in parallel with the functional specialisation of distinct plastid types (Choi et al. 2021; Bohnsack and Schleiff 2010; Kessler and Schnell 2006). Accordingly, multiple TOC component isoforms assemble into functionally-distinct complexes that preferentially mediate the import of either photosynthetic or non-photosynthetic proteins; these are broadly referred to as photosynthetic (TOC-P) and non-photosynthetic (TOC-N) pathways (Andrès et al. 2010; Nellaepalli et al. 2026). Core TOC components are the β-barrel channel Toc75 and two GTPase receptors of low (∼30 kDa) and high (∼100-150 kDa) molecular weight (Sun and Jarvis 2023; Shi and Theg 2013a; Kessler and Schnell 2006); these receptors exist in multiple isoforms, respectively forming the Toc33 and Toc159 receptor families. Both receptor classes contain a GTPase (G) domain and a transmembrane helix (TMH), while high-molecular-weight receptors additionally possess a membrane-embedded β-barrel M domain. In the Toc159 family, the Toc159 protein primarily mediates the import of photosynthetic proteins, whereas Toc120 and Toc132 act redundantly in the import of mainly non-photosynthetic preproteins (Ivanova et al. 2004; Kubis et al. 2004). The Toc159 family receptors have an N-terminal acidic (A) domain that plays an important role in determining client recognition specificity (Agne et al. 2009; Ivanova et al. 2004; Inoue et al. 2010).

While the functional specialisation of TOC receptors during early plastid biogenesis is well established, accumulating evidence indicates that the composition of the TOC machinery is dynamically regulated across tissues and developmental stages (Kubis et al. 2004; Infanger et al. 2011a; Ling et al. 2012, 2019, 2021a; Nellaepalli et al. 2026). The channel Toc75 is rapidly upregulated and assembled at the onset of plastid development, followed by sequential recruitment of members of the Toc159 and Toc33 receptor families, which increase in abundance as differentiation progresses (Nellaepalli et al. 2026). Such observations suggest that plastid protein import is not static but instead undergoes coordinated remodelling during development. During senescence, TOC abundance is also controlled by CHLORAD-mediated turnover (Ling et al. 2012, 2019, 2021a). However, CHLORAD is a TOC-removal system rather than an active participant in the import mechanism, and whether individual GTPase receptors actively regulate later plastid maturation and leaf ageing remains unclear.

Among the Toc159 receptor family members, Toc90 is the least characterised. Early studies reported physical interaction of Toc90 with Toc75 and Toc33, and suggested a potential genetic interaction with Toc159 (Hiltbrunner et al. 2004). Furthermore, overexpression of Toc90 in a *toc159* background was reported to partially rescue the mutant’s albino phenotype (Infanger et al. 2011a). However, the physiological relevance of these findings was unclear as constitutive overexpression only delivered partial rescue. Consistent with this uncertainty, *toc90* loss-of-function mutants do not display a discernible phenotype under standard growth conditions (Kubis et al. 2004; Ramundo et al. 2020). The evolutionary origin of Toc90 also remains ambiguous. While some studies have proposed Toc90 to be an ancestral member of the Toc159 receptor family, others have misleadingly assigned the Toc90 name to homologous proteins within the Toc120 clade, highlighting inconsistencies in its classification (Liu et al. 2023; Jin et al. 2022). In the absence of clear information on its function and evolutionary roots, Toc90 has often been regarded as an ancient form of the Toc159-type receptor (Hofmann and Theg 2003; Reumann et al. 2005).

Here, we resolve the function and evolutionary origin of Toc90 using a combination of genetic, physiological, biochemical and structural approaches. We show that Toc90 is a *bona fide* TOC receptor and define its functional specificity. Furthermore, phylogenetic analyses reveal that Toc90 is not an ancestral receptor, but rather a recently evolved member of the Toc159 family, consistent with a model of neofunctionalisation within the Toc159 receptor group. Together, our findings uncover a previously unrecognised layer of plastid protein import regulation, in which the TOC machinery is dynamically remodelled during chloroplast ageing.

## Results

### Toc90 manipulation alters translocon component stoichiometry but does not impair chloroplast development

To assess the biological relevance of Toc90, we began by examining whether Toc90 plays a measurable phenotypic role *in vivo*. We generated independent transgenic overexpression lines expressing the full-length *TOC90* CDS under the control of the *35S* CaMV promoter in the wild-type background. The resulting overexpression lines (OE) accumulated approximately threefold higher levels of Toc90 protein compared to wild type (**Supplementary Fig. 1A; Supplementary Table ST1**). In parallel, we examined previously-described *toc90* T-DNA insertion knockout (KO) lines, *toc90-1* and *toc90-3* (Kubis et al. 2004). However, neither the KO nor the OE lines were distinguishable from wild type with respect to overall growth phenotype, chlorophyll content, or maximal photosynthetic efficiency at 12-15 days after germination (DAG) **(Fig. 1A**, **Supplementary Fig. 1B, C)**.

**Figure 1.**
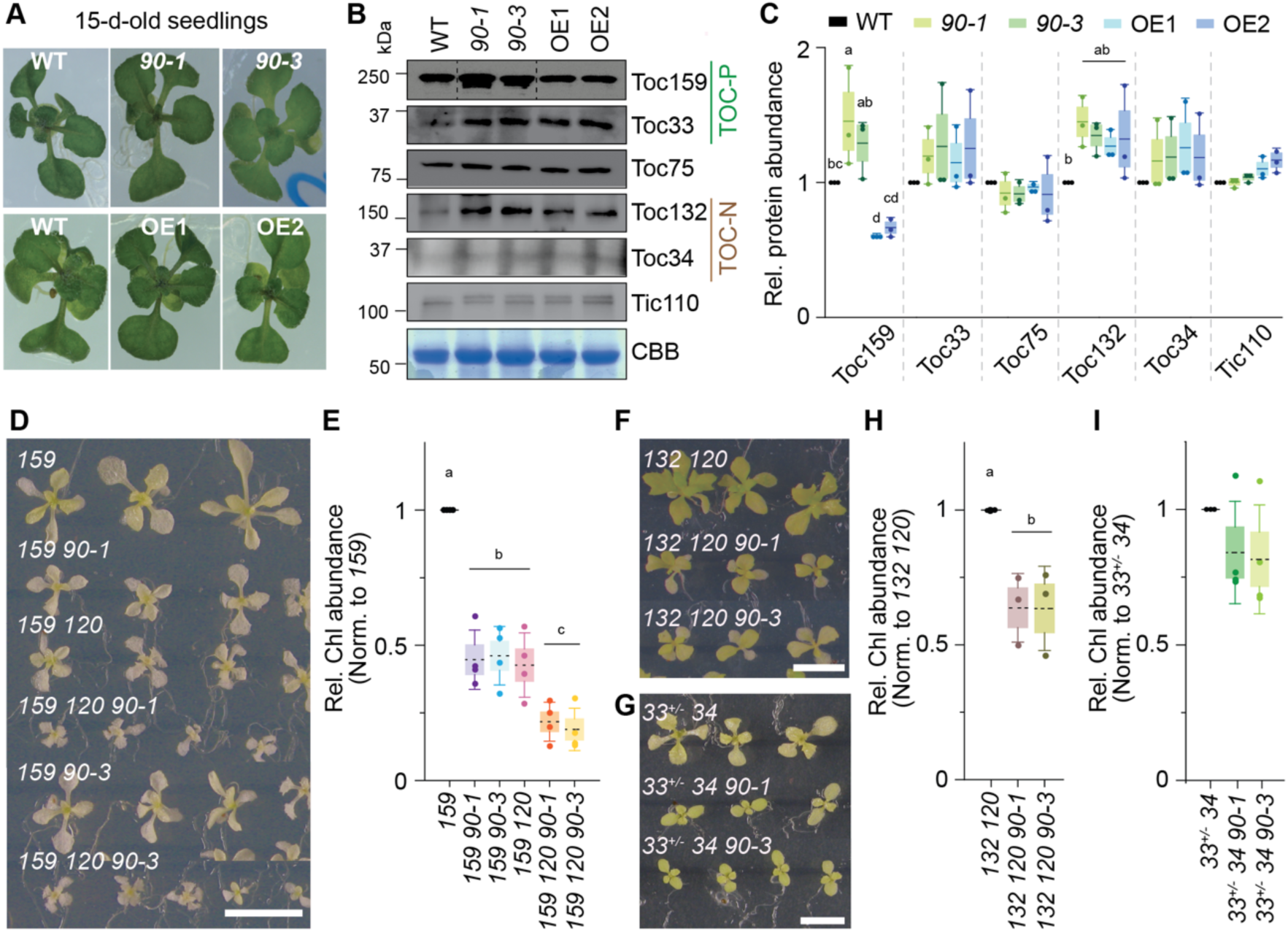
Phenotypic and molecular characterisation of AtToc90. **(A)** Phenotypic images of 15-day-old seedlings of WT, *90-1*, *90-3*, OE1 and OE2 grown on half-strength MS medium, captured using a Zeiss dissecting microscope (10× magnification), Scale bar = 0.2 cm. **(B)** Relative protein levels of the chloroplast translocon components Toc159, Toc33, Toc75, Toc132, Toc34, and TIC110 were analysed by immunoblotting using specific antibodies. Total protein extracts were prepared from equal amounts of fresh tissue harvested from 15-day-old seedlings. Coomassie Brilliant Blue (CBB) staining of a parallel gel is shown as a loading control. **(C)** Densitometric quantitation of the above-mentioned TIC-TOC components from at least three independent biological replicates. Every blot intensity is normalised to total lane intensity recorded from the loading control gel. **(D-I)** Phenotypes and relative chlorophyll content of double and triple mutant lines generated by crossing *toc90-1* or *toc90-3* into the parental genotypes (D) *toc159* and *toc159 toc120* and (F) *toc120 toc132* and (G) *toc33^+/-^ toc34* (Scale bar = 0.5 cm). Seedlings were initially grown for 12 days on half-strength MS medium and then transferred to medium containing 0.5% sucrose. For panels (D) and (F) (top), images were taken 32 days after seedling emergence; for panel G (bottom), images were taken 24 days after emergence. Chlorophyll content was measured prior to sucrose transfer using DMF extraction from 10-12-days-old seedlings. Statistical analysis in (C, E, H and I) was performed using one-way ANOVA followed by Fisher’s least significant difference (LSD) test for pairwise comparisons among genotypes. Homogeneity of variance was assessed using Levene’s test.

When equal amounts of tissue were analysed, the accumulation levels of core TOC-TIC components, including Toc75, Toc33, Toc34 and Tic110, were not changed significantly in the KO and OE lines **(Fig. 1B, C)**; Toc33 and Toc34 showed a slight increase in protein abundance, but these differences were not statistically significant. In contrast, the two large GTPase receptors exhibited differential accumulation patterns: Toc132 levels were increased by approximately 30-50% in both KO and OE lines compared to wild type, whereas Toc159 displayed reciprocal changes, with its abundance increased by ∼25-40% in KO lines but decreased by ∼25-30% in OE lines **(Fig. 1B, C)**. These differences suggest compensatory adjustments within the TOC-TIC complex in response to changes in Toc90 abundance, most likely to maintain the complex stoichiometry of TOC components. Moreover, the inverse relationship between Toc90 and Toc159 levels indicate coordinated regulation between these receptor isoforms.

### Genetic interactions reveal limited functional overlap between Toc90 and canonical receptors

Given that Toc90 manipulation altered the abundance of other TOC components, we further investigated the functional links between the components by generating a comprehensive series of double- and triple-knockout mutant lines. The *toc90-1* and *toc90-3* alleles were crossed into previously-established single-mutant backgrounds (*toc159, toc132*, *toc120, toc75-III, toc75-IV, toc33* and *toc34*), as well as into double-mutant backgrounds (*toc159 toc120, toc132 toc120^+/-,^ toc132 toc120, toc132 toc33^+/-^, toc132 toc33* and *toc33^+/-^ toc34*) **(Fig. 1D-I; Supplementary Figs. 2, 3)**.

With the exception of *toc159 toc90-1* and *toc159 toc90-3* **(Fig. 1D-I)**, none of the double mutants exhibited detectable differences in phenotype or chlorophyll accumulation relative to their respective parental lines **(Supplementary Figs. 2, 3)**. Toc159 *toc90* double mutants displayed pronounced defects, accumulating approximately 60-65% less chlorophyll than the *toc159* single mutant. These effects were further exacerbated in triple-mutant backgrounds. The *toc159 toc120 toc90* triple mutants were markedly smaller than the *toc159 toc120* parent and accumulated 50-55% less chlorophyll; relative to *toc159* alone, chlorophyll levels were reduced by 75-80% **(Fig. 1E)**. Similarly, homozygous *toc132 toc120 toc90* plants were smaller and accumulated ∼35% less chlorophyll compared to the *toc132 toc120* parent line **(Fig. 1F, H)**.

Similar observations were made with the heterozygous mutant genotypes **(Fig. 1F-I)**. The *toc132 toc120^+/-^ toc90* triple mutants were smaller and accumulated ∼20-28% less chlorophyll than the respective parental genotype. While the t*oc34 toc33^+/-^ toc90* triple mutants did appear smaller than parental genotype plants, the differences in chlorophyll were not statistically significant. The reduced impact of Toc90 manipulation in the latter background may reflect the fact that Toc33/34 are of a different receptor family and is consistent with the unchanged accumulation of Toc33 and Toc34 observed in *toc90* single-mutant backgrounds.

Taken together, these results demonstrate that Toc90 is functionally relevant and influences chloroplast protein import, as reflected by reduced plant size and chlorophyll content in specific mutant combinations. The relatively modest (∼25-35%) reductions in chlorophyll caused by loss of Toc90 in the *toc132 toc120* background suggest only limited redundancy between Toc90 and the TOC-N receptor components. In contrast, the severe (50-75%) chlorophyll loss caused by loss of Toc90 in the *toc159* background, together the compensatory changes in Toc159 accumulation seen in the Toc90 KO and OE lines, strongly supports a closer functional association between Toc90 and the Toc159-dependent (TOC-P) import pathway. However, unlike the reciprocal complementation observed between Toc120 and Toc132 in the corresponding single mutants, TOC90 does not compensate for the *toc159* albino phenotype (Kubis et al. 2004; Infanger et al. 2011a). This suggests two non-mutually exclusive possibilities: (1) Toc90 is unlikely to serve as a principal receptor for photosynthetic preproteins and instead provides limited functional backup when Toc159 activity is constrained; and/or (2) the substantially lower native expression of Toc90 relative to Toc159 restricts its compensatory capacity in *toc159* mutant backgrounds.

### Toc90 possesses an acidic domain that confers partial functional redundancy with Toc159

To address the two potential explanations for the limited functional overlap between Toc159 and Toc90, we carried out a series of *in silico* and genetic analyses. While the G- and M-domains of Toc90 are well structured, its N-terminal region is intrinsically disordered, consistent with the A-domains of other Toc159-family receptors **(Supplementary Fig. 4A)**. Historically studies have proposed either no A-domain or very short non-acidic N terminal extension for Toc90 (Kubis et al. 2004; Infanger et al. 2011a; Hiltbrunner et al. 2004; Infanger et al. 2011b; Bischof et al. 2011; Shi and Theg 2013b), but NCBI CDD domain searches suggest an N-terminal extension toward Glu-135 **(Supplementary Fig. 4B)**. Using modern structural prediction approaches, including the AlphaFold model of Toc90 **(Supplementary Fig. 4C, D)** (Abramson et al. 2024), we observed low-confidence, disordered structure up to Lys-88, followed by a clear transition to ordered α-helical structure after residue 89 **(Fig. 2A)**. This was supported by changes in the phi and psi dihedral angles to values characteristic of the geometry of an α-helix **(Supplementary Fig. 4E)**. Disorder predictions (DISOPRED3) further supported this boundary, indicating strong intrinsic disorder restricted to residues 1-83, a property also presents among other the other family members **(Supplementary Fig. 4F)**. Biochemical features were consistent with this assignment of the Toc90 A-domain: residues 1-83 exhibited a low isoelectric point (pI) and strong enrichment of acidic residues (a characteristic feature of the A-domains), while extending the region of analysis to residue 135 increased the pI and reduced the charge asymmetry **(Fig. 2B)**. Together, these analyses indicate that the definition of the Toc90 A-domain needs to be revised, and that a more appropriate mapping spanning residues 1-83 should be considered.

**Figure 2.**
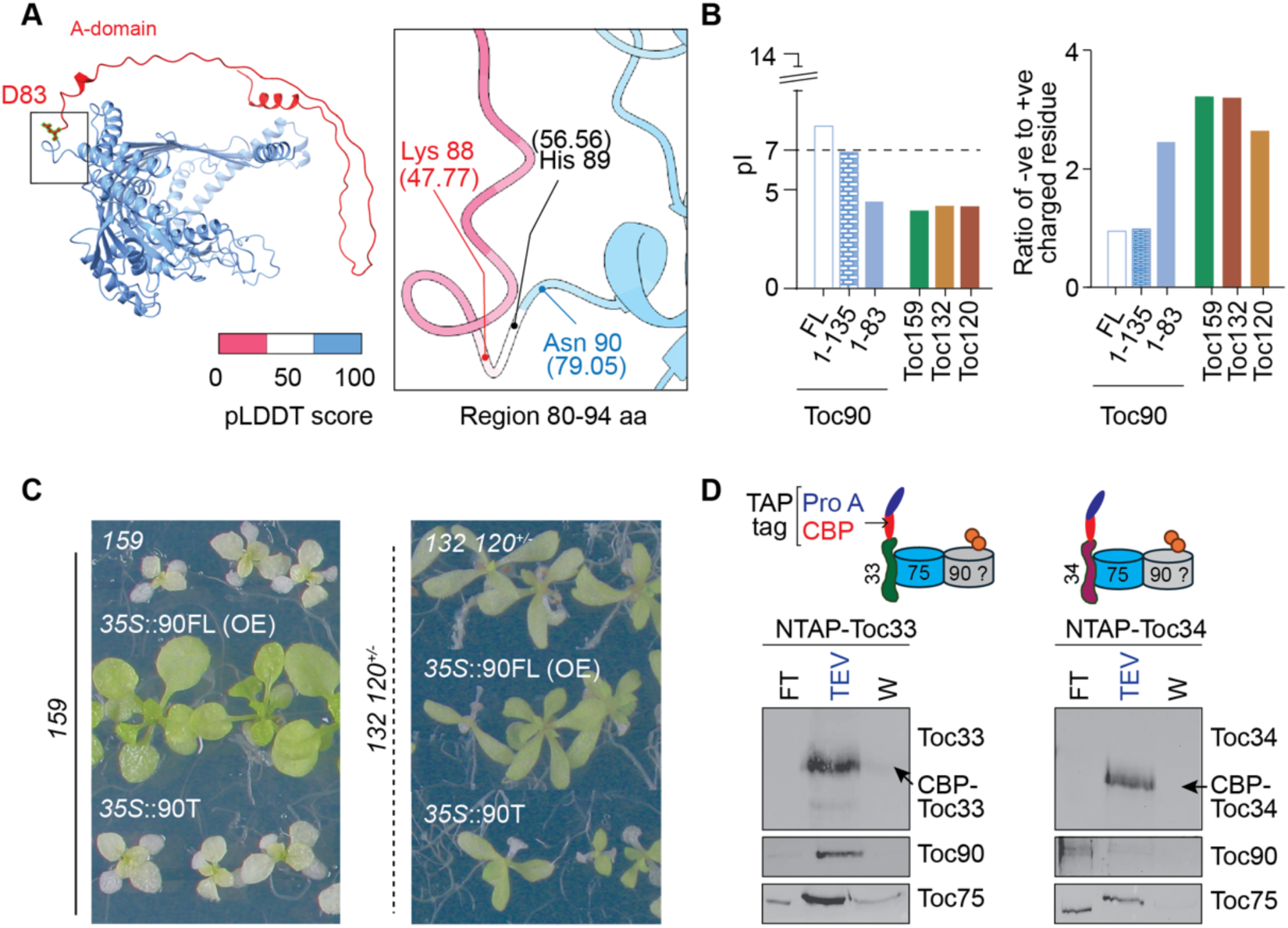
Role of acidic domain of Toc90 and Toc90’s associating partners. **(A)** AlphaFold-predicted structure of full-length AtToc90 and the corresponding per-residue confidence scores (pLDDT). Higher pLDDT values indicate greater confidence in the predicted structure. The inset highlights amino acids 80-94, showing a marked increase in pLDDT values following Lys88, suggesting improved local structural confidence in this region. **(B)** Comparison of isoelectric points (pI) among Toc159 family members. pI values are shown for full-length AtToc90 (FL), its N-terminal 1-135 amino acid region, and the 1–83 amino acid region. These values are compared with the acidic domains of other Toc159 family members: AtToc159, AtToc120, and AtToc132. The ratio of positively to negatively charged residues in full-length AtToc90 and its truncated regions is also compared with that of the corresponding acidic domains of the other Toc159 family proteins. **(C)** Phenotypic analysis of AtToc90 overexpression lines in toc159 and toc132 toc120^+/-^ mutant backgrounds. Full-length AtToc90 CDS (*35S*::90FL same as OE) and truncated AtToc90 CDS (*35S*::90T) were expressed in the indicated backgrounds. Seedlings were grown for 12 days on half-strength MS medium and then transferred to medium containing 0.5% sucrose and imaged 24 days after emergence. **(D)** AtToc90 is physically associated with photosynthetic TOC complexes. Schematic representation of NTAP-*Toc33* (photosynthetic TOC) and NTAP-*Toc34* (non-photosynthetic TOC) complexes, in which Protein A (ProtA) and calmodulin-binding peptide (CBP) tags are fused to the N terminus of Toc33 and Toc34, respectively. Immunoblot analysis of Toc33, Toc90, and Toc75 in NTAP-Toc33-purified complexes, and of Toc34, Toc90, and Toc75 in NTAP-Toc34 - purified complexes. Three fractions: post-binding flow-through (FT), TEV-eluted fraction, and SDS wash (W) were collected, separated by SDS–PAGE, and probed with the indicated antibodies.

Kubis et al. (2004) reported that *TOC90* transcript levels are more than 60% lower than those of *TOC159*, suggesting that *TOC90* may not reach the expression threshold required to sustain the import in young, rapidly developing tissues normally mediated by Toc159. To test whether increased expression could compensate for this limitation, we overexpressed different Toc90 coding sequences in the *toc159* KO background. Based on evidence that the A-domain of Toc159 family proteins is important for preprotein recognition specificity (Kessler and Schnell 2006; Inoue et al. 2010), we focused on two Toc90 variants expressed under the *35S* CaMV promoter: a full-length construct (*35S*::90FL; this is the same as the OE construct), and a truncation lacking 14 N-terminal residues (*35S*::90T; this was analogous to the construct used by (Kubis et al. 2004)). Under identical growth conditions, the *35S*::90FL partially rescued the albino *toc159* phenotype as expected (Infanger et al. 2011a), whereas the truncated variant delivered no detectable complementation effect **(Fig. 2C)**. Overexpression of both constructs was confirmed at both transcript and protein levels, and chlorophyll levels were determined to verify the complementation results **(Supplementary Fig. 5)**.

These findings demonstrated that a complete Toc90 A-domain is essential for the receptor’s limited functional redundancy with Toc159. Moreover, even when expressed under a strong constitutive expression, Toc90 can only partially substitute for Toc159, indicating that the import of Toc159-dependent preproteins is likely not its primary physiological role.

We next assessed potential redundancy with the TOC-N receptors by overexpressing both constructs in the *toc132 toc120^+/-^* background. Neither *35S*::90FL nor *35S*::90T altered the phenotype, supporting the view that Toc90 does not functionally overlap with Toc120 or Toc132. Accordingly, the ∼30% chlorophyll reduction observed in *toc132 toc120 toc90* triple mutants, relative to the respective control, likely reflects a cumulative “load effect” caused by simultaneous disruption of multiple import pathways, rather than genuine functional redundancy. This contrasts with the obvious functional redundancy seen in the Toc159-Toc90 relationship.

Collectively, these results suggest a specific functional connection between Toc90 and Toc159, with partial redundancy mediated by the acidic A-domain of Toc90.

### Toc90 physically interacts with TOC-P complex components

Given the genetic evidence supporting a functional connection between Toc90 and TOC-P, we next investigated whether Toc90 can physically interact with the TOC-P components. To this end, we performed affinity purifications using transgenic lines expressing TAP-tagged Toc33 (a TOC-P component) or Toc34 (a TOC-N component) **(Supplementary Results 1)**. Purifications were performed using IgG Sepharose matrix (which binds the protein A domain of the TAP tag), and serial elutions performed using TEV protease and SDS (Wash -W) were analysed alongside post-binding flowthrough (FT) by immunoblotting **(Fig. 2D)**.

In NTAP-Toc33 purifications, full-length TAP-Toc33 was largely retained on the IgG column and efficiently eluted by TEV cleavage, with ∼65% recovered in the TEV fraction (estimated by densitometric quantification of the Toc33 signal across the FT, TEV and W fractions). The Toc75 protein co-purified strongly with Toc33 as expected. Interestingly, more than 80% of Toc90 was present in the TEV eluate, indicating that Toc90 is associated with Toc33-containing complexes **(Fig. 2D)**. In NTAP-Toc34 purifications, similar enrichment of TAP-Toc34 in the TEV fraction was seen, but Toc75 recovery was lower, consistent with the reduced abundance of Toc34 in greening tissue (Nellaepalli et al. 2026). Importantly, >70% of Toc90 was detected in the FT fraction in this case, rather than in the TEV eluate as seen with Toc33, indicating that Toc90 associates only weakly with Toc34-containing complexes. It is important to note that, because antibody sensitivities vary, the relative intensities of different proteins on the blots cannot be compared directly. Nevertheless, these results clearly establish that Toc90 preferentially associates with Toc33-containing TOC complexes and not with the Toc34-containing TOC complexes.

### TOC90 emerged as a distinct seed-plant lineage within the TOC159 receptor family

The results described above show that Toc90 shares functional similarity with Toc159 and is physically associated with components of TOC-P complexes. Therefore, we wished to assess the extent to which Toc90 shares ancestry with Toc159. A previous phylogenetic study, though limited in scope, indicated that Toc90 is basal to the Toc159 family clade(Kubis et al. 2004). Much later, two Toc159 family members present in green algae were named Toc90 and Toc120 on the basis of limited sequence analysis (Jin et al. 2022; Liu et al. 2023; Ramundo et al. 2020). Overall, such observations tended to suggest that Toc90 may represent an ancient, early-evolving ancestor of the higher-plant Toc159 family (Hofmann and Theg 2003; Reumann et al. 2005).

To clarify the evolutionary relationship between Toc90 and the other members of the Toc159 family, we constructed a phylogenetic tree using 262 curated Toc GTPase receptor sequences from 67 species spanning green algae, bryophytes, ferns, gymnosperms and angiosperms **(Fig. 3; Supplementary Fig. 6)**. Only sequences containing both the G-domain and the M-domain were included. As the N-terminal A-domain is highly variable and difficult to align reliably across divergent taxa, the phylogeny was reconstructed using only the more conserved G- and M-domain regions.

**Figure 3.**
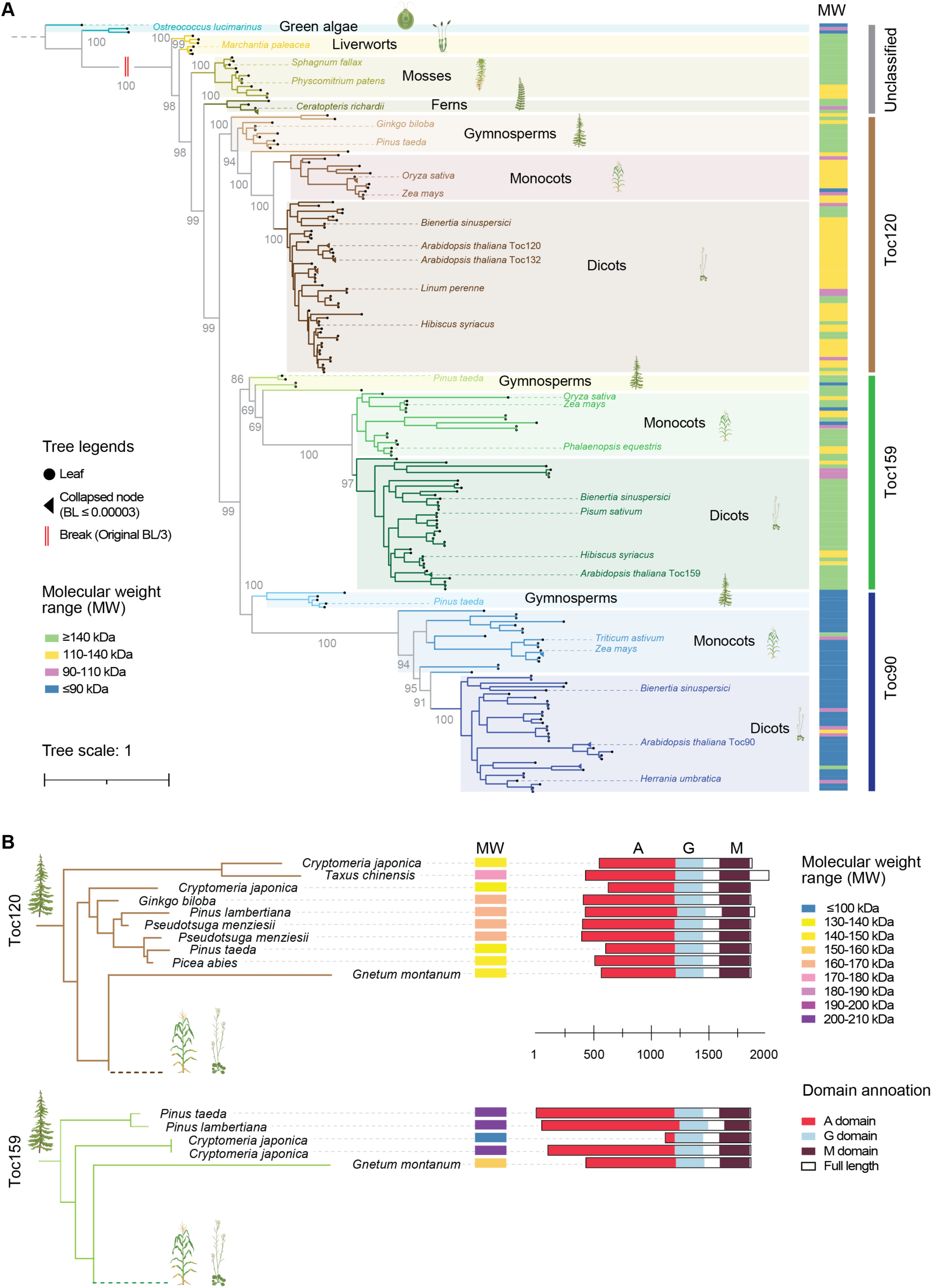
Evolutionary diversification of Toc159-family receptors and the emergence of Toc90. **(A)** Phylogenetic tree of Toc159-family proteins from green algae, bryophytes, ferns, gymnosperms, monocots, and dicots. The analysis resolves three major seed-plant receptor lineages corresponding to Toc90, Toc159, and Toc120 represented in blue, green and brown lines; while sequences from green algae and early-diverging land plants remain unclassified and shown in grey lines at the rightmost side of the tree. Branch colours also correspond to the major receptor groups: Toc120, Toc159, Toc90, and unclassified groups. Background shading marks major plant lineages. Black circles indicate terminal leaves, black triangles indicate collapsed nodes with very short branch lengths (<0.0003), and red perpendicular lines between green algae and liverworts indicate a break introduced in long branches for visualisation. Numbers at selected nodes indicate branch support bootstrap (1000) values in percentage. The molecular-weight strip on the right shows the predicted size class of each protein, highlighting that Toc90 sequences are predominantly shorter than Toc120 and Toc159 members. Branch lengths represent sequence divergence, with longer branches indicating greater accumulated amino-acid substitutions. Well-known species are mentioned in their respective positions in the tree, full tree with all species name and bootstrap value are provided in **Supplementary figure 6**. **(B)** Phylogenetic placement and domain organisation of representative gymnosperm Toc159-family and Toc120-family proteins. Predicted molecular-weight classes are shown beside each sequence, and domain schematics indicate the predictive positions of the acidic A-domain, GTPase G-domain, and membrane-associated M-domain as obtained from NCBI CDD search.

The resulting tree broadly followed the expected trajectory of plant evolution **(Fig. 3A)**. Green algal, liverwort, moss, and fern sequences were positioned outside or near the base of the three canonical seed-plant receptor clades and were therefore classified separately. These early-diverging lineages contained Toc GTPase receptors with heterogeneous molecular weights. However, the smaller proteins from these lineages did not cluster within the seed-plant Toc90 clade. Thus, low molecular weight alone is not sufficient to define a Toc90 orthologue. Instead, these early-diverging proteins are better interpreted as ancestral or unclassified Toc159-family receptors. This indicates that Toc90 is unlikely to represent an ancient receptor form inherited directly from algae.

Within seed plants, Toc159-family receptors formed three clades that we named Toc120, Toc159 and Toc90 after the archetypal Arabidopsis proteins. Each clade contains gymnosperm, monocot and dicot sequences, which suggests that the three lineages diverged before the major seed-plant groups separated. The corresponding Toc120 and Toc159 proteins are predominantly of high molecular weight, consistent with the retention of large N-terminal A-domains; in contrast, the Toc90 proteins are generally below 110 kDa. Toc90 therefore appears to have evolved through marked A-domain reduction while retaining the conserved G-M receptor core.

In gymnosperms, Toc120-type proteins were already smaller than Toc159-type proteins, which shows that Toc120 and Toc159 diverged as distinct full-length receptor lineages before angiosperm evolution rather than through recent shortening of Toc159 **(Fig. 3B)**. Further size reductions in both groups likely occurred within angiosperms.

Branch lengths increased in the order Toc120<Toc159<Toc90, particularly in angiosperms **(Fig. 3A)**. This suggests stronger conservation of the Toc120-type proteins, consistent with a constrained housekeeping role. This observation is also consistent with the hypothesis that the Toc159-type proteins underwent specialisation for photosynthetic protein import. The even more pronounced divergence of Toc90 suggests the acquisition of new functions. In fact, the long branches and reduced size of angiosperm Toc90-type proteins support its emergence as a derived, potentially neofunctionalised receptor. Although small (∼100 kDa) proteins were present before the gymnosperm-angiosperm split, their strong separation from angiosperm Toc90 sequences suggests that the major functional specialisation of true Toc90 protein occurred during angiosperm evolution.

### The Toc90 complex is globally similar to the Toc159 containing TOC-P complex but displays distinct local structural features

Having established that Toc90 is a neofunctionalised paralogue of Toc159 that has undergone at least partial functional divergence, we next asked whether its structural features had similarly diverged during angiosperm evolution. As Toc90 associates with Toc33 and Toc75 **(Fig. 2D)**, we used AlphaFold 3 (AF3) to model the putative Toc33-Toc75-Toc90 complex (Toc90 complex) **(Fig. 4A; Supplementary Fig. 7)**. To do this, we used the mature sequence of theToc75, the G- and M-domains of Toc90, full-length Toc33, two GTP molecules, and two Mg²⁺ ions **(Fig. 4A; Supplementary Fig. 7A)**. The multimeric complex was well supported by an interface predicted template modelling (iPTM) score of 0.76, predicted template modelling (pTM) score of 0.74, and an overall predicted local distance difference test (pLDDT) score of 78.1, and there was excellent structural confidence at potential regions of interaction **(Supplementary Fig. 7)**.

**Figure 4.**
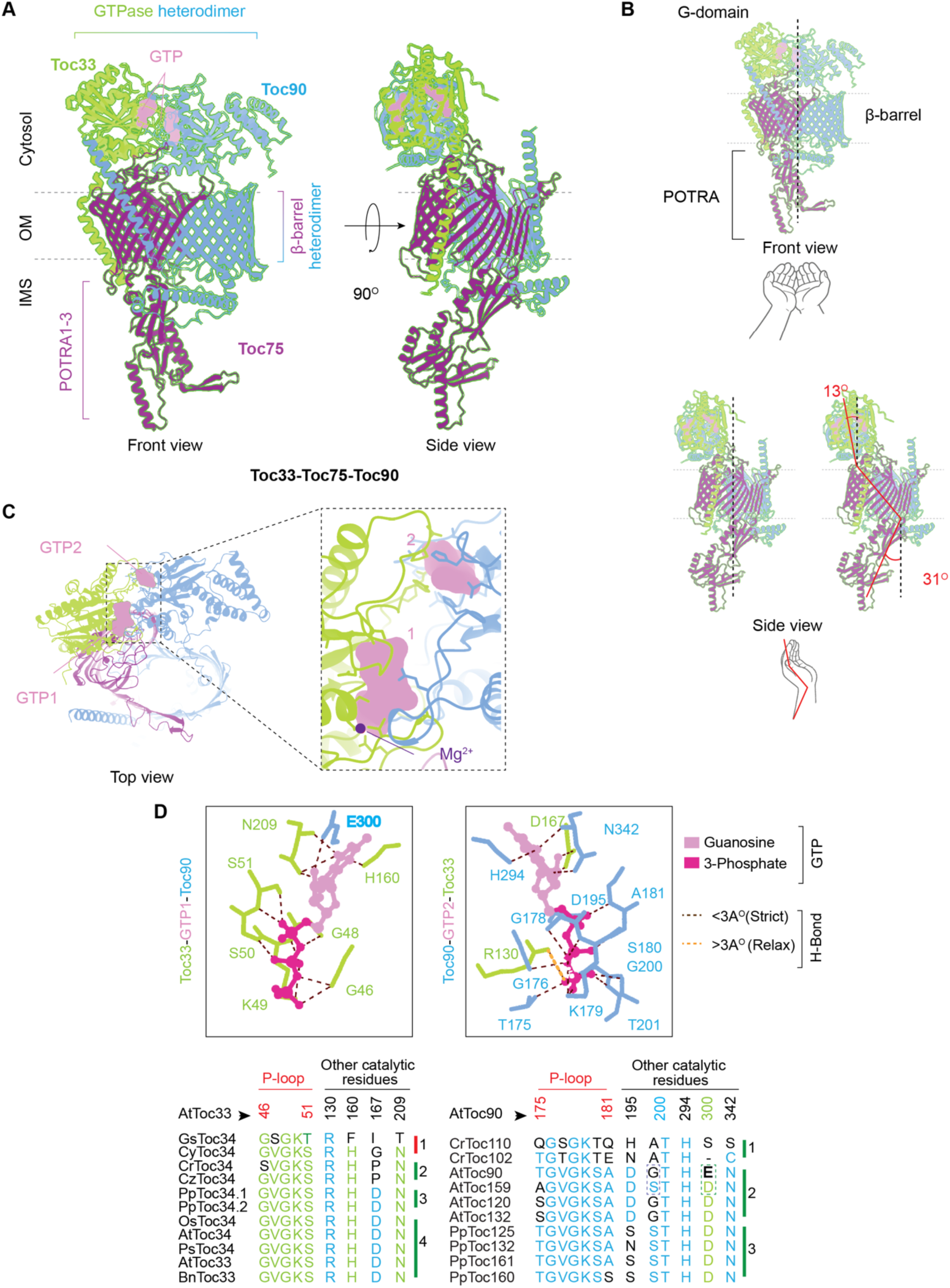
AF3-predicted architecture of the Toc33-Toc75-Toc90 complex and conservation of interfacial GTP-binding sites. **(A)** AlphaFold3-predicted structural model of the Toc33-Toc75-Toc90 complex shown in front and side views. Toc33 is shown in green, Toc75 in magenta, Toc90 in cornflower blue, and bound GTP in pink. The approximate membrane position is indicated with cytosol, outer membrane (OM), and intermembrane space (IMS) labels. Toc75 forms a β-barrel in the outer membrane and extends into the IMS through POTRA1–3 domains, whereas Toc33 and Toc90 form a cytosolic GTPase heterodimer. The side view represents a 90° rotation around the y axis. **(B)** Relative arrangement of the AF3-predicted Toc33-Toc75-Toc90 heterotrimer. Front, side, and top views show the orientation of the G domains and the positioning of the two nucleotide-binding sites, GTP1 and GTP2. The inset highlights the close arrangement of the two GTP-binding pockets, including the Mg²⁺-coordinated nucleotide. Angular deviations indicated in red summarise the relative tilt between the compared GTPase domains and the POTRA domains compared to the central axis of the complex. **(C)** Close-up views of the two predicted nucleotide-binding pockets at the Toc33-Toc90 interface. Toc33 residues are shown in green, with neighbouring Toc90 residues in blue. Guanosine and triphosphate moieties are shown in light and dark pink, respectively. Hydrogen bonds are indicated by dashed lines, with black denoting strict contacts below 3Å and orange denoting relaxed contacts above 3Å. Sequence alignments below each pocket show conservation of key nucleotide-contacting residues across representative Toc33/Toc34 and Toc90/Toc159 homologs. At - *Arabidopsis thaliana*; Gs - *Galdieria sulphuraria*; Cy - *Chlamydomonas yellowstonensis*; Cr - *Chlamydomonas reinhardtii*; Cz - *Chlamydomonas zimbabwiensis*; Pp - *Physcomitrium patens*; Os - *Oryza sativa*; Bn - *Brassica napus*.

The Toc90 polypeptide retains the characteristic G-M architecture **(Supplementary Fig. 8A)** as previously noted for Toc159 (Nellaepalli et al. 2026). The Toc90 G-domain possesses the conserved GTPase fold comprised of eight α-helices (named H1-H8) and a half β-barrel comprised of six β-strands (named β1-β6) **(Supplementary Fig. 8B-D)**; and it possesses the five highly-conserved nucleotide-binding motifs (boxes G1-G5), including the phosphate-binding P-loop (G1 box) which secures the GDP/GTP molecule **(Supplementary Fig. 8C)**. The Toc90 M-domain comprises a three-helix connector region, a 14-stranded β-barrel, and a transmembrane helix (TMH) **(Supplementary Fig. 8A,E)**. The long unfolded loop at the C-terminal end of the connector polypeptide remains embedded within the Toc75 β-barrel as previously noted for Toc159 (Nellaepalli et al. 2026). Thus, overall, despite its sequence divergence, Toc90 retains the principal structural features expected for a Toc159-family GTPase receptor.

The predicted Toc90 complex adopts a cupped-hand appearance, in which the GTPase heterodimer forms a finger-like cytosolic module, the β-barrel heterodimer forms a cupped palm region, and the POTRA domains extend towards the intermembrane space like a forearm **(Fig. 4B; Supplementary Fig. 9A,B)**. In the front view of the GTPase heterodimer, the two GTP molecules (GTP1 in the Toc33 catalytic pocket; GTP2 in the Toc90 catalytic pocket) are positioned close to the central axis of the complex, whereas in the top view GTP1 lies towards the front of the GTPase heterodimer and GTP2 lies behind it; each nucleotide is associated with a nearby Mg²⁺ ion **(Fig. 4C, inset)**. Both GTP molecules predominantly interact with P-loop and catalytic elements of their cognate GTPase while making more limited contacts with the opposing receptor **(Fig. 4D)**.

We subsequently compared the Toc90 model with the previously reported Toc33-Toc75-Toc159 (TOC-P) complex (Nellaepalli et al. 2026). Overall, the two complexes are highly similar, with most aligned regions showing RMSD values below 5 Å **(Supplementary Fig. 9C;)**. The GTPase heterodimer and the β-barrel heterodimer were both similar to those reported previously for the TOC-P complex (Nellaepalli et al. 2026) **(Supplementary Fig. 9C)** (Nellaepalli et al. 2026). Thus, the predicted Toc90 complex preserves the overall architecture of the canonical Toc159-containing TOC-P complex. Despite this global similarity, six local regions (marked a-f) differed between the Toc90 and Toc159 complexes, including the nucleotide-binding pocket and the intermembrane-space-facing linker region **(Supplementary Fig. 9C)**. In particular, the bound GTP molecules adopted different predicted orientations **(Supplementary Fig. 9D)**, while the Toc90 nucleotide-binding pocket displayed a more negative electrostatic environment and altered hydrophobicity compared with Toc159 **(Supplementary Fig. 9E)**. Notably, GTP1 interacts with Toc90 residue E300, whereas the corresponding position is occupied by aspartate (D) in other Toc159-family proteins in *Arabidopsis* and in other species **(Fig. 4D)**. Although this D-to-E substitution preserves the negative charge, the additional methylene (-CH_2_-) group of glutamate (E) may have subtly altered the local geometry of the nucleotide-binding pocket by shifting the position of the terminal carboxylate group. GTP2, by contrast, interacts with Toc33 residues R130 and D167, which are broadly conserved among Toc33/Toc34 homologues, including red-algal representatives **(Fig. 4D)**. These changes could also influence the catalytic rate of the GTPases imparting functional implications.

Together, the altered nucleotide orientation, nucleotide-pocket electrostatic properties, and organisation of the linker region reveal specific local structural differences between the otherwise strongly conserved architecture of the Toc90 and TOC-P complexes. These differences may influence receptor interactions, conformational flexibility, and/or coupling between the G- and M-domains. Thus, they provide a structural basis through which Toc90 may have acquired a specialised biological role.

### Toc90 expresses specifically at later stages of development

TOC receptor components are known to display tissue- and developmental-stage-specific expression patterns (Kubis et al. 2004; Constan et al. 2004; Kubis et al. 2003; Dahlin and Cline 1991; Teng et al. 2012). We previously found that Toc90 abundance remained largely unchanged during the first three weeks after seedling emergence, in contrast to its close homologue Toc159, which is highly expressed during early development (Dahlin and Cline 1991; Kubis et al. 2004; Nellaepalli et al. 2026). We therefore examined later developmental stages to identify conditions under which Toc90 abundance might change.

As an initial assessment, we performed a pilot glasshouse experiment comparing the development of wild type with the *toc90* and OE lines. No obvious morphological differences were observed among the genotypes **(Supplementary Fig. 10A)**. We also compared bolting time, recorded as the number of weeks after germination, as an indicator of accelerated or delayed development. However, neither mutant nor OE lines showed a detectable change in bolting time relative to the wild type **(Supplementary Fig. 10B)**.

We next examined naturally occurring changes in Toc90 abundance in wild-type plants during later development **(Fig. 5A)**. Toc90 abundance increased markedly in seven-week-old plants reaching more than twice the level detected at three weeks. In contrast, the abundance of all other TOC complex components declined over the same period. Toc159 showed reduction by ∼70% in seven-week-old plants relative to three-week-old plants. By comparison, Toc132 was retained to a substantially greater extent, though still showed a ∼30% reduction. Both Toc33 and Toc34 decreased by ∼40%, whereas Toc75 declined by ∼20%. The inner-envelope translocon component Tic110 was also reduced, by approximately ∼30% **(Fig. 5A)**. Since the absolute stoichiometry of TOC components is uncertain, relative abundances cannot be used to define the precise composition of the complexes. Nevertheless, the pronounced reduction in Toc159, together with the comparatively smaller reduction in Toc33 and Toc75 and the more than twofold increase in Toc90, raises the possibility that Toc90-containing complexes become proportionally more abundant during later development.

**Figure 5.**
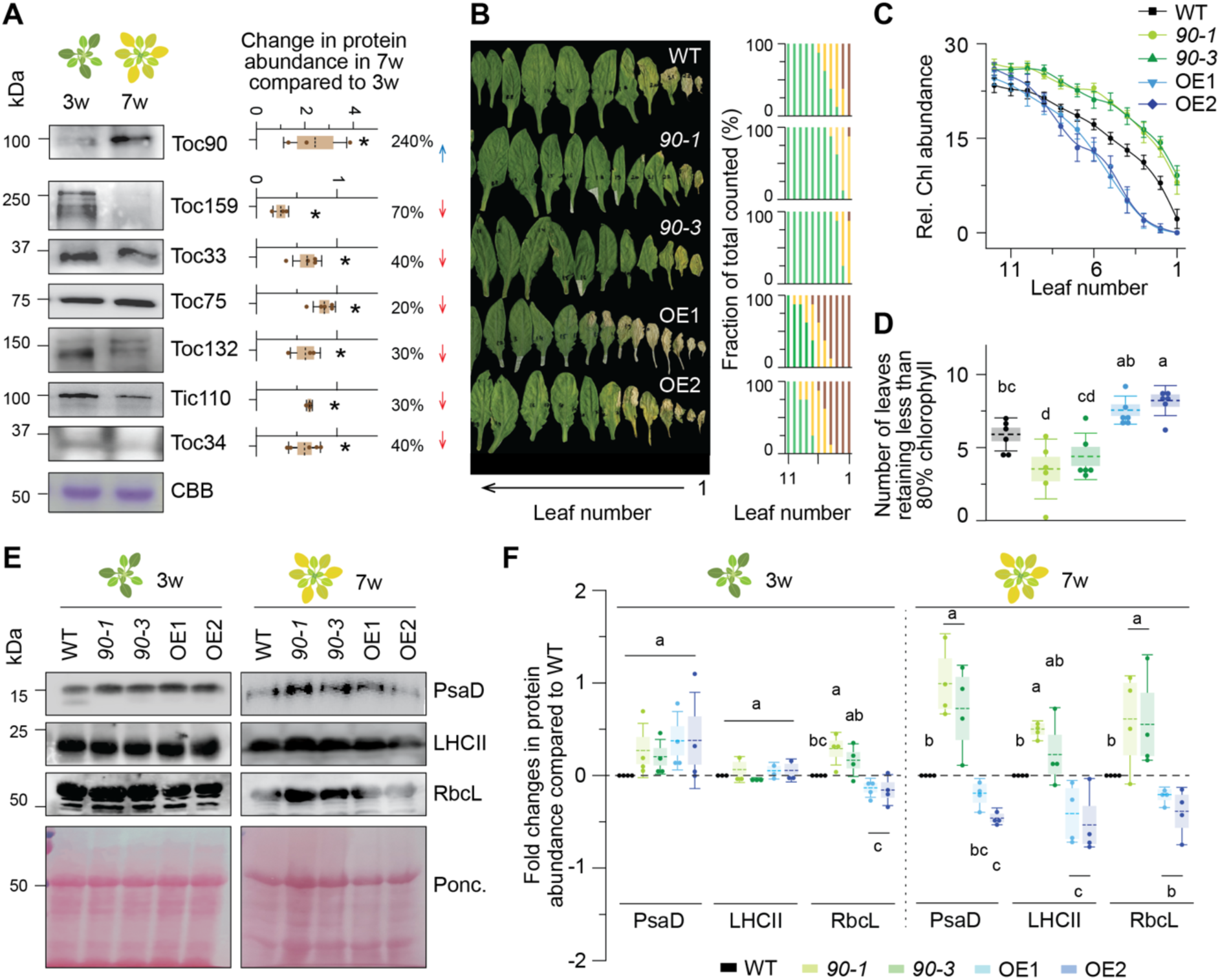
Role of Toc90 in leaf ageing. **(A)** Immunoblot analysis of different chloroplast translocon proteins in 3-week-old and 7-week-old Arabidopsis rosettes. Protein levels were quantified from immunoblots and normalised to total protein loading (30 µg per lane), shown by Coomassie Brilliant Blue (CBB) staining. Densitometric quantification on the right shows relative percentage changes in protein abundance in 7-week-old plants compared with 3-week-old plants (n=3); Asterisks indicate significance level p<0.05. Corresponding red down or blue up arrow indicate down or upregulation. **(B)** Leaf-by-leaf visual senescence phenotype of WT, two *toc90* mutant lines, and two Toc90 overexpression lines. Leaves were arranged from older to younger leaves according to leaf number. *toc90* mutant plants retained greener older leaves, whereas Toc90 overexpression lines showed stronger chlorosis in older and middle-aged leaves. Corresponding graphs indicate visual scoring of leaves in three categories, green (non-senescent), yellow (senescing) and brown (dead) (n=8). **(C)** Quantification of relative chlorophyll abundance across successive leaves measured using a SPAD-502 meter (Konica-Minolta) (n=5-9). For each biological replicate at least values of 5 plants have been averaged. **(D)** Number of leaves with less than 80% relative chlorophyll content compared to greenest leaves (Average of leaf number 12-15 per plants were averaged), a gold standard for leaf senescence onset. The chlorophyll depletion rates were determined by fitting the SPAD values with a Boltzmann function. **(E)** Immunoblot analysis of representative photosynthetic proteins in wild type, *toc90 KO*, and OE lines at 3 and 7 weeks. PsaD, LHCII, and RbcL were used as markers for photosystem I, light-harvesting complex II, and Rubisco/chloroplast protein abundance, respectively. Ponceau staining was used as a loading control. **(F)** Quantification of PsaD, LHCII, and RbcL abundance relative to wild type at 3 and 7 weeks (n=3-4). Since in 7 week *toc90* lines show higher retention of proteins (mainly RbcL and LHCII) we refrained from normalising the densitometric quantitation based on total protein per lane. Individual points represent biological replicates, box indicates SE, and error bars show SD among replicates. Statistical analysis in (A, D and F) was performed using one-way ANOVA followed by Fisher’s least significant difference (LSD) test for pairwise comparisons among genotypes. Homogeneity of variance was assessed using Levene’s test. Different letters denote statistically significant differences among genotypes in F.

It is conceivable that a subset of Toc33-Toc75 complexes (Nellaepalli et al. 2026) may associate with Toc90 as Toc159 abundance declines. The comparative retention of Toc132 and Toc34 may indicate that Toc132-Toc34-containing complexes are maintained independently. This would be consistent with our evidence that Toc90 preferentially associates with Toc33 rather than Toc34 **(Fig. 2D)**. Together, these results reveal a developmental shift in the relative abundance of TOC receptor components and suggest that Toc90 may acquire greater functional importance during later stages of plant development.

### Toc90 promotes age-dependent developmental senescence

As Toc90 abundance increases markedly around seven weeks, we revisited our pilot glasshouse experiment to determine whether altered Toc90 abundance was associated with a late developmental phenotype. Interestingly, the OE plants appeared to senesce earlier than both wild-type and *toc90* KO plants **(Supplementary Fig. 10A)**. Consistent with this observation, chlorophyll abundance was lower in the OE and wild-type plants than in KO plants, suggesting that loss of Toc90 may delay age-related senescence **(Supplementary Fig. 10A)**.

To examine this phenotype more systematically, we performed a developmental senescence assay using 49-day-old plants **(Fig. 5B; Supplementary Fig. 11)**. The total number of rosette leaves did not differ among the lines, indicating that the observed differences were not caused by major changes in leaf production or developmental timing. To quantify senescence progression, individual leaves were visually classified into three categories: green, non-senescent leaves; yellow, senescing leaves; and brown, fully senescent leaves **(Fig. 5B)**. For leaf 1, 10% of individuals were brown for KO compared to 100% for both wild-type and OE plants. For leaf 6, none of the individuals were yellow for KO compared to ∼15% for wild type and 100% for OE plants. Finally, for leaf 10, none of the individuals were yellow for KO and wild-type plants compared to ∼15% for OE plants. Thus, senescence had progressed in substantially younger leaves in the OE lines than in the other genotypes.

The visible phenotypes described above were consistent with measurements of chlorophyll abundance **(Fig. 5C)**. A reduction to below ∼80% of the chlorophyll level of non-senescent leaves is commonly associated with the onset of senescence (Lihavainen et al. 2023, 2020; Fataftah et al. 2021). On average, approximately 6 leaves per wild-type plant, 4-5 leaves per KO plant, and 7-9 leaves per OE plant contained less than 80% chlorophyll **(Fig. 5D)**. Together, these findings indicate that Toc90 positively contributes to age-dependent developmental senescence and may promote the onset or progression of senescence-associated chlorophyll degradation.

### Loss of Toc90 delays senescence-associated degradation of photosynthetic proteins

Because Toc90 functions as a chloroplast protein-import receptor, we hypothesised that the delayed senescence of *toc90* KO plants might result from altered chloroplast protein homeostasis. We therefore examined whether altered Toc90 abundance affects the developmental decline of representative photosynthetic proteins **(Fig. 5E)**. To this end, we compared photosynthetic protein abundance in three- and seven-week-old plants. The reason for selecting these two ages is related to the fact that the accumulation of the photosynthetic proteins occurs during early developmental stages (Kubis et al. 2003; Nellaepalli et al. 2026), whereas at later stages such proteins are generally maintained at static levels (Nelson and Millar 2015). Thus, comparing the two was expected to reveal any developmental effects of Toc90 manipulation on photosynthetic protein components, due to the partial functional redundancy between Toc159 and Toc90.

We analysed a core component of photosystem I (PsaD), a major photosynthetic antenna complex protein light-harvesting complex II (LHCII), and the Calvin–Benson-cycle enzyme ribulose-1,5-bisphosphate carboxylase/oxygenase (RuBisCO). In three-week-old plants, PsaD and LHCII abundance did not differ significantly between the wild-type, KO and OE plants, although PsaD showed a modest increase in both transgenic backgrounds. RuBisCO abundance followed a pattern similar to that observed for Toc159, being slightly higher in the KO lines and lower in the OE lines relative to wild type **(Fig. 5E,F)**. Consistent with the observed visible phenotypes, at seven weeks the abundance of PsaD, LHCII and RuBisCO had declined markedly in the wild-type and OE plants but notably remained substantially higher in the KO plants. Compared to wild type, the KO lines retained ∼40-50% more PsaD and RuBisCO and ∼25% more LHCII, relative to the earlier time point. By contrast, the OE lines retained ∼25-50% less of each protein than the wild type **(Fig. 5E,F)**.

Together, these findings suggest that Toc90 positively contributes to the dismantling of the photosynthetic apparatus during age-dependent senescence. As a chloroplast import receptor, Toc90 may facilitate the import of factors required for chlorophyll breakdown, photosynthetic-complex disassembly, or stromal protein degradation.

### Toc90 promotes stress-induced senescence

Having established that Toc90 contributes to age-dependent developmental senescence, we next asked whether it also influences senescence induced by external stress signals. We first examined dark-induced leaf senescence by covering individual mature leaves to exclude light (Weaver and Amasino 2001). After eight days of darkness, leaves from both *toc90* KO lines remained visibly greener than those from wild-type plants. Consistent with their visible appearance, the covered KO leaves retained significantly more chlorophyll than the corresponding wild-type leaves **(Supplementary Fig. 12A,B)**.

We next investigated whether Toc90 also affects senescence-like responses induced by oxidative stress (Ling and Jarvis 2015). Seven-to ten-day-old seedlings were transferred to medium containing paraquat, a herbicide that specifically targets the photosynthetic electron transport chain to induce ROS production and oxidative stress. After 11 days, the paraquat treatment induced pronounced leaf yellowing and chlorophyll loss in the wild-type plants. Under the same conditions, both *toc90* KO lines remained visibly greener and retained a greater proportion of green leaves than the wild type **(Supplementary Fig. 12C,D)**.

Together, these findings support a broader role for Toc90 in regulating the onset or progression of senescence. Given its function as a chloroplast protein-import receptor, Toc90 may facilitate the import of chloroplast proteins required for stress-induced chlorophyll degradation, photosynthetic-complex disassembly, or associated retrograde signalling.

## Discussion

Chloroplast protein import has been studied primarily in the context of organelle biogenesis when rapidly developing plastids require the high-capacity import of numerous photosynthetic and metabolic proteins (Choi et al. 2021; Bohnsack and Schleiff 2010; Kessler and Schnell 2006). The expression of most TOC components is high during such early developmental stages, but in later stages when biogenesis slows down the accumulation of TOC proteins is reduced (Jarvis et al. 1998; Kubis et al. 2003; Ling and Jarvis 2015; Nellaepalli et al. 2026; Dahlin and Cline 1991; Chu and Li 2018) **(Fig. 6A)**. In this context, it is understandable that the lack of obvious phenotypic defects in seedlings lacking Toc90 was taken to indicate that Toc90 is a non-functional protein. Here, we correct this misconception by identifying Toc90 as a neofunctionalised Toc159-family receptor whose principal physiological contribution becomes apparent during the later stages of development.

**Figure 6.**
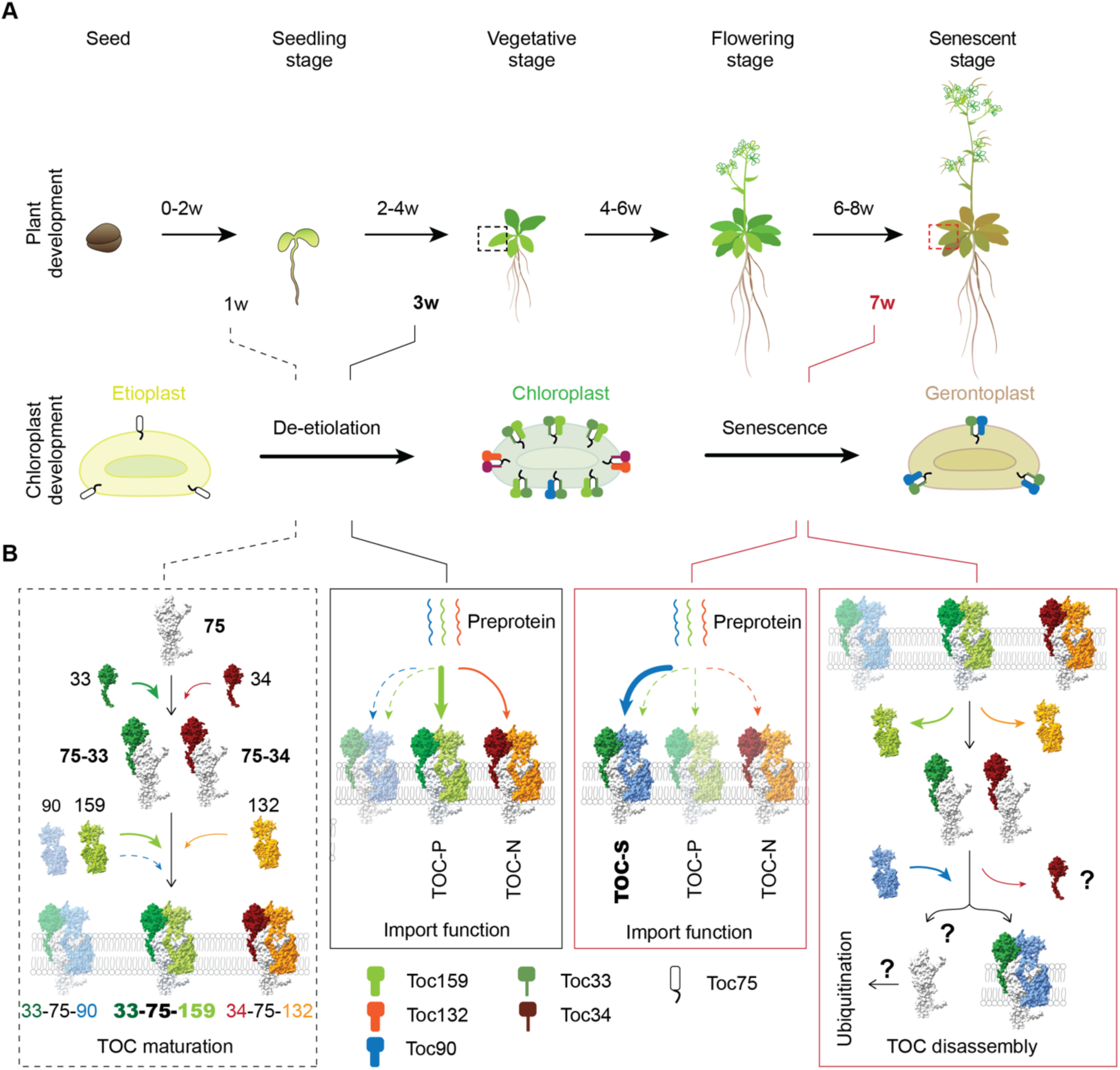
Working model for TOC translocon dynamics across plant development. Schematic representation of the Arabidopsis developmental cycle, from seedling establishment to the reproductive stage, together with the concomitant progression of plastid/chloroplast development from the etioplast stage. **(A)** During early chloroplast development, approximately one to three weeks after germination, TOC-P and TOC-N complexes predominate and mediate the import of photosynthetic and housekeeping precursor proteins. At these stages, Toc90 abundance remains low relative to Toc159- and Toc132-containing complexes. **(B)** At later developmental stages, particularly during the transition from flowering to reproductive growth, Toc90 accumulates, whereas Toc159 and Toc132 levels decline. Based on the data presented here, Toc90 may become incorporated into Toc75-Toc33-containing complexes, forming a Toc90-Toc75-Toc33 translocon : **TOC-S**. The pathway responsible for degradation or turnover of the Toc90-containing complex remains unclear. Dashed box indicates data adopted only from the literature. The black box indicates a conclusion based on both experimental data from this paper and previous literature and the red box indicates a conclusion based on experimental evidence in this manuscript.

### Toc90 is a functional and specialised member of the Toc159 receptor family

The lack of a strong early phenotypic defects in *toc90* mutants previously suggested that Toc90 may be vestigial or largely redundant (Kubis et al. 2004; Reumann et al. 2005; Jarvis 2008). However, our data support a developmentally conditional role for Toc90 rather than the absence of an important function. Altered Toc90 abundance affected the levels of other large TOC receptors, particularly Toc159 and Toc132, suggesting coordinated regulation within the TOC receptor network **(Fig. 1)**, potentially through compensatory expression or competition for complex stoichiometry. Genetic and biochemical evidence further supported such possibilities: full-length Toc90 partially complemented *toc159* mutant plants, whereas an N-terminally truncated form did not; while Toc90 preferentially co-purified with Toc33 rather than Toc34. These findings indicated that Toc90 acts within a Toc33-dependent pathway similar to that mediated by TOC-P but is not functionally interchangeable with Toc159 **(Fig. 2)**. Moreover, phylogenetic analysis supported the notion that Toc90 is a relatively recently derived Toc159 paralogue in seed plants, rather than an ancestral receptor with algal origins. Reduction or loss of ancestral protein regions following gene duplication is a recognized route to paralogue diversification and, in some cases, functional specialisation or neofunctionalisation (Weiner 3rd et al. 2006; Casewell et al. 2011; Force et al. 1999; Man et al. 2020) **(Fig. 3)**. The compact architecture and pronounced divergence of Toc90 in angiosperms are therefore consistent with such an evolutionary trajectory. In agreement with these ideas, AF3 modelling predicted that the Toc33-Toc75-Toc90 complex retains the overall architecture of the Toc159-containing TOC-P complex (Nellaepalli et al. 2026), while showing local differences around the GTP-binding pockets and G-M connector **(Fig. 4)**. These distinctive features may influence nucleotide cycling, receptor communication, and/or substrate selectivity, providing a structural basis for the specialised role of Toc90.

### Toc90 promotes and modulates senescence

The Toc90 protein accumulated post-bolting, at around seven weeks of age with the onset of senescence, as Toc159 and other TOC-TIC components declined **(Fig. 5A)**. This suggested that chloroplast ageing involves remodelling of receptor composition rather than a uniform loss of import capacity. Thus, Toc90 may provide a late-stage pathway adapted to the import requirements of senescing chloroplasts. This would complement well the established role of ubiquitin-dependent control in TOC machinery regulation (Sun and Jarvis 2023), wherein SP1 and the chloroplast-associated protein degradation (CHLORAD) pathway promote the removal of TOC components to modulate import and enable plastid proteome reconfiguration during developmental transitions and environmental stress (Ling et al. 2021b, 2019, 2017; Ling and Jarvis 2016, 2013). Such CHLORAD-mediated regulation also influences the timing of leaf senescence, demonstrating that control of the import apparatus is directly connected to plastid ageing (Sun et al. 2022; Mohd Ali et al. 2023). The increase in Toc90 may represent the complementary side of this remodelling process: while biogenesis-associated receptors are selectively removed or reduced, Toc90 may be retained, stabilised or newly produced to support (perhaps in conjunction with newly available Toc75 and/or Toc33) residual import programmes required for chloroplast dismantling **(Fig. 6A,B)**.

The senescence phenotypes associated with Toc90 manipulation point to the existence of a late-stage import programme. Because rosette leaf number and bolting time were largely unchanged, the delayed yellowing in *toc90* was not attributable to slower vegetative development **(Supplementary Fig. 10B)**. Instead, senescence progressed more slowly along the leaf-age gradient in *toc90* KO mutants, and was extended into younger leaves in Toc90 OE plants **(Fig. 5B-D)**, indicating that Toc90 abundance influences senescence timing. Consistently, seven-week-old *toc90* KO plants retained higher levels of PsaD, LHCII and RbcL, while these proteins declined more strongly in the OE lines **(Fig. 5E, F)**. The coordinated retention of these proteins in the mutant indicates broad preservation of the photosynthetic apparatus and delayed chloroplast dismantling. The *toc90* KO mutants also showed reduced senescence responses to darkness and paraquat treatment, indicating that Toc90 contributes to a pathway shared by developmental and stress-induced chloroplast degradation. Thus, Toc90 may import components of a common senescence-execution programme (Hörtensteiner 2006; Woo et al. 2019; Matile et al. 1999). Alternatively, loss of Toc90 may preserve a photosynthetic or redox state that raises the threshold for senescence initiation (Krieger-Liszkay et al. 2019; Mayta et al. 2019).

### Potential mechanistic models for Toc90 action during senescence

Based on our results, one can envision at least two different models for the role of Toc90 during senescence, both of which involve a special senescence-associated TOC complex termed TOC-S (comprised of Toc90, Toc33 and Toc75). In the first of these models (direct cargo model), Toc90 preferentially recognises a subset of nucleus-encoded proteins required for chlorophyll catabolism, photosystem disassembly, proteolysis, and/or redox signalling. Candidate Toc90 client classes would include chlorophyll-catabolic enzymes and their regulators, chloroplast proteases and protease adaptors, thylakoid-remodelling proteins and components that communicate chloroplast redox status to the nucleus (Hörtensteiner 2006; Woo et al. 2019; Matile et al. 1999). In the absence of Toc90, insufficient accumulation of one or more such factors would delay degradation of chlorophyll and photosynthetic proteins, and thereby retard the whole developmental programme **(Fig. 6B)**.

In the second model (complex composition substitution model), Toc90 does not recognise a narrow class of “senescence enzymes,” but instead changes the overall selectivity or capacity of the residual TOC apparatus. Incorporation of Toc90 could displace Toc159, alter Toc33 activity, or change the conformational properties of the Toc75 channel. This could shift the balance of imported proteins away from photosynthetic maintenance and towards chloroplast dismantling. It is conceivable that displaced and unattached Toc159 would then become an easier target for ubiquitination and CHLORAD-mediated degradation. In this model, the mutant phenotype would arise from the cumulative effects of altered import across multiple chloroplast proteins, leading to a broader shift in chloroplast proteome composition, rather than from the failed import of a single critical cargo **(Fig. 6B)**.

In conclusion, our findings identify Toc90 as a neofunctionalised Toc159-family receptor that exhibits an expression profile biased towards late development, and which links chloroplast protein import with leaf senescence. We propose that chloroplast ageing involves coordinated remodelling of the TOC machinery, in which declining Toc159-dependent biogenesis pathways are accompanied by an increased contribution of Toc90-containing (TOC-S) complexes. The delayed chlorophyll loss and prolonged retention of photosynthetic proteins in *toc90* KO mutants signifies the identification of a novel stay-green phenotype (Thomas and Ougham 2014; Gregersen et al. 2013), highlighting Toc90 as a potential target for manipulating senescence and chloroplast longevity in crops.

## Methods

### Plant material and growth conditions

All wild-type, mutant and transgenic plant lines used in this study were *Arabidopsis thaliana* ecotype Columbia-0. The two *toc90* knockout mutants used in this study were *toc90-1* (1236-C11) and *toc90-3* (NG19434) (Hiltbrunner et al. 2004; Infanger et al. 2011a; Kubis et al. 2004), and have been described previously. Genotypes were verified by PCR using gene-specific and T-DNA border primers listed in Supplementary Table S1. Other TOC mutants used in this study have been described previously (Kubis et al. 2004; Constan et al. 2004; Jarvis et al. 1998; Kubis et al. 2003).

Seeds were surface sterilised and grown on half-strength Murashige and Skoog (½ MS) medium (pH 7.5) containing vitamins supplemented with 0.7% agar, with or without 0.5% (w/v) glucose as indicated, using previously described procedures (Hiltbrunner et al. 2004; Infanger et al. 2011a; Kubis et al. 2004). Plants used for soil-grown experiments were either transferred to soil after 10-12 d of growth on ½ MS medium or sown directly on soil. Directly sown seeds were stratified for 3 days at 5 °C before transfer to the growth cabinet. Unless otherwise stated, plants were grown under long-day conditions (16 h light/8 h dark) at 20 °C, 100-120 µmol photons m⁻² s⁻¹ and approximately 60% relative humidity. For germination assays, surface-sterilised seeds were sown on MS agar plates, stratified at 5 °C for 3 days and subsequently maintained in a Percival Scientific growth chamber at 20 °C under long-day conditions at 100-120 µmol photons m⁻² s⁻¹.

### Transgenic line generation

For generation of AtToc90-overexpressing lines, the *AtTOC90* coding sequence was amplified by PCR using primers Toc90 (H4C12)-F (5′-CATCTGGCATACCGGAGCTTGA-3′) and Toc90 (H4C12)-R (5′-CTTCGAAACACGGCGGGACTT-3′). The amplified product was first cloned into the Gateway entry vector pDONR207 and subsequently transferred by LR recombination into the binary destination vector pB2GW7 (Karimi et al. 2002). pB2GW7 carries spectinomycin resistance for bacterial selection and a phosphinothricin/Basta resistance marker for plant selection. The resulting construct was verified by restriction analysis using SacI and XhoI. Wild type plants were transformed with the above construct via *Agrobacterium* with floral dip method. Toc34 tandem affinity purification (TAP) fusion constructs were generated using the Gateway-compatible plant vectors pNTAPi and pCTAPi (Rohila et al. 2004). The *TOC34* coding sequence was amplified using primers listed in Supplementary Table 1; all additional primers used in this study are listed in Supplementary Table 2

### Physiological experiments

In vivo leaf chlorophyll abundance was estimated using a Konica-Minolta SPAD-502 meter as described previously. Chlorophyll was also quantified after extraction in either N,N-dimethylformamide (DMF) (Moran 1982) or 80% acetone (Porra et al. 1989). For chlorophyll measurements, 3-8 independent experiments (biological replication) were performed; approximately 9-15 seedlings or 3-5 leaves from different plants were pooled per genotype in each experiment. Maximum quantum efficiency of photosystem II (Fv/Fm) was determined using a CF Imager (Technologica) after 30 min dark incubation as described previously (Ling et al. 2012), with 3-5 seedlings analysed per experiment.

### Protein extraction and immunodetection

Total proteins were extracted from equal fresh weights of seedlings or leaves as described previously (Kovacheva et al. 2005). Protein concentrations were determined by the Bradford assay using the Bio-Rad Protein Assay Kit (Bradford 1976). Samples were normalised either to fresh weight or total protein content, as appropriate for the experiment, and separated by SDS-PAGE using 10% polyacrylamide gels for Toc159 and Toc132 or 12% gels for all other proteins. Following electrophoresis, gels were either stained with Coomassie brilliant blue or proteins were transferred to polyvinylidene difluoride (PVDF) membranes by wet transfer. Membranes were blocked in Tris-buffered saline containing 0.1% Tween 20 (TBST; pH 7.5) supplemented with 2% blocking-grade milk for detection of Tic110, Toc90, Toc75, Toc34, Toc33, PsaD and LHCII; 5% blocking-grade milk for Toc159 and RbcL; or 10% blocking-grade milk for Toc132. Subsequent immunodetection was performed as described previously (N. Li and Jarvis 2024).

TOC and TIC components and chloroplast marker proteins were detected using antibodies against the AtToc75-III POTRA domain (1:1,000), AtToc159 A-domain (1:5,000), AtToc132 A-domain (1:1,000), an AtToc33 peptide (1:500), Toc34 (1:2,000; AS07 238, Agrisera), the AtTic110 stromal domain (1:5,000), PsaD (1:10,000; AS09 461, Agrisera), LHCII (1:5,000; described previously (Dall’Osto et al. 2017)) and RbcL (1:2,000; described previously (Havaux et al. 2007)). The AtToc90 antibody was raised against the synthetic peptide SDPFFRDPHQEQDNHC (Eurogentec). An anti-rabbit IgG antibody conjugated to horseradish peroxidase was used as the secondary antibody (1:5,000; 12-348, Sigma-Aldrich). Chemiluminescence was detected using the EZ-ECL Enhanced Chemiluminescence Detection Kit for HRP (Biological Industries, Sartorius) and an ImageQuant LAS 4000 imaging system (GE Healthcare). Band intensities were quantified using ImageJ version 1.54u8 (Schneider et al. 2012). Densitometric analyses were based on at least three independent experiments showing consistent trends; representative immunoblots are shown in the figures.

### Phylogenetic analysis

For phylogenetic analysis, a curated sequence dataset was assembled from the National Center for Biotechnology Information (NCBI), Phytozome, The Arabidopsis Information Resource (TAIR) and PlantGenIE. Conserved domains and diagnostic motifs in retrieved proteins were examined using the NCBI Conserved Domain Database (CDD) (Lu et al. 2020), and domain-validated sequences were used as queries for additional homology searches. Taxonomic sampling included dicots, monocots, ferns, gymnosperms, mosses, clubmosses, liverworts, green algae, rhodophytes, cyanobacteria and proteobacteria. Candidate homologues were identified using BLASTp (Altschul et al. 1990) with an E-value threshold of 0.05 and a maximum of 250 target sequences per search. Searches used the BLOSUM45 substitution matrix, gap costs of 13 for opening and 3 for extension, a word size of 5 and a maximum of two matches per query range. Conditional compositional score-matrix adjustment was enabled, and no additional low-complexity filtering or masking was applied during the initial search phase.

The highly disordered and clade-variable N-terminal acidic (A) domains were identified using DISOPRED3 (Jones and Cozzetto 2015) and removed before multiple-sequence alignment, thereby focusing the analysis on the conserved central GTPase (G) and C-terminal membrane-associated (M) domains.

Amino acid sequences were aligned using MAFFT version 7 with the E-INS-i iterative refinement strategy(‘MAFFT Multiple Sequence Alignment Software Version 7: Improvements in Performance and Usability | Molecular Biology and Evolution | Oxford Academic’, n.d.). Maximum-likelihood phylogenies were inferred using IQ-TREE version 3.0.1 (Wong et al. 2026). The best-fitting amino acid substitution model was selected by ModelFinder (Kalyaanamoorthy et al. 2017) according to the Bayesian Information Criterion; Q.PLANT+R7 was selected for the Toc159-family dataset. Q.PLANT is a plant-specific empirical amino acid replacement matrix (Minh et al. 2021), while +R7 models among-site rate heterogeneity using seven FreeRate categories. Branch support was assessed using 1,000 ultrafast bootstrap replicates (UFBoot2) (Hoang et al. 2018). The resulting consensus tree was visualised and annotated using Interactive Tree of Life (iTOL) (Letunic and Bork 2024).

### AlphaFold 3 structural modelling and analysis

Three-dimensional structural models of TOC receptor-channel complexes were generated using the AlphaFold 3 Server (https://alphafoldserver.com) (Abramson et al. 2024). Multimeric receptor-channel assemblies were modelled to investigate structural organisation, protein-protein interfaces and potential functional specialisation among TOC receptor components.

For modelling Toc159- and Toc90-containing complexes, the following components were provided as input: (i) the mature Toc75 sequence (D141-Y818); (ii) Toc159 lacking its N-terminal acidic (A) domain (E781-Y1503) or the corresponding A-domain-truncated Toc90 sequence (L84-S793); (iii) full-length Toc33 (M1-L297); (iv) two GTP molecules; and (v) two Mg²⁺ ions. Five structural predictions were generated for each receptor complex and ranked using the confidence metrics provided by AlphaFold 3. All predictions were inspected, and the highest-ranked model for each complex was used for detailed structural analysis and figure preparation.

Model confidence and predicted protein-protein interfaces were evaluated using predicted alignment error (PAE), predicted local distance difference test (pLDDT), predicted template modelling (pTM) and interface predicted template modelling (ipTM) scores. Structural visualisation, superposition and interface analysis were performed using UCSF ChimeraX (Pettersen et al. 2021). Molecular surfaces were coloured according to Coulombic electrostatic potential in ChimeraX, and residue hydrophobicity was represented using the Kyte-Doolittle scale (Kyte and Doolittle 1982).

### Senescence assays

Natural leaf senescence was assessed by allowing plants to develop under standard growth conditions for up to 7 weeks. Rosette leaves were numbered sequentially according to their order of emergence, with the oldest rosette leaf designated as leaf 1, consistent with established Arabidopsis leaf-senescence analyses. At the onset of visible bolting, defined by elongation of the inflorescence stem, the youngest visible rosette leaf was marked by introducing a small hole in the leaf blade. This mark served as a positional reference for subsequent identification of individual rosette leaves as plants continued to develop. Plants were photographed at the indicated developmental stages to document whole-plant senescence. At 7 weeks, rosette leaves were detached and arranged sequentially from the oldest to the youngest leaf to visualise the progression of natural senescence across leaf positions (Yang et al. 2014).

For individual leaf dark-induced senescence assays, 9^th^ rosette leaves from 4 week old plants were maintained in darkness for 5 days while remaining attached to the intact plant, following the principle of the individually darkened leaf assay. To minimise mechanical damage and permit gas exchange, each selected leaf was first enclosed in a small sleeve prepared from a single layer of breathable cotton fabric and shaped to accommodate the leaf. The sleeve was subsequently covered with aluminium foil to exclude light completely. The remainder of the plant was maintained under normal growth conditions. After 5 days of dark treatment, the covers were removed and leaves were photographed and analysed for senescence-associated phenotypes (Weaver and Amasino 2001).

For chemically induced senescence, seedlings were germinated on ½ MS medium containing vitamins and lacking glucose and grown for 14 days after emergence. Individual seedlings were then transferred to ½ MS agar plates supplemented with 10 µM paraquat and maintained under the same growth conditions for 11 days. Senescence-associated phenotypes were monitored during the treatment, and seedlings were photographed after 11 days of paraquat exposure.

### Statistical analysis

Statistical analyses were performed using Origin 8.1 (OriginLab). Data are presented as individual raw data points together with the mean ± standard error (SE). Statistical significance was assessed by one-way analysis of variance (ANOVA). When significant differences among groups were detected, pairwise comparisons were performed using either Tukey’s post hoc test or Fisher’s least significant difference (LSD) test, as indicated in the corresponding figure legends. Homogeneity of variances was assessed using Levene’s test. Exact *P* values or significance thresholds are provided in the corresponding figure legends. For multiple-group comparisons, groups labelled with different letters were considered significantly different.

