## Supplementary information for "Neofunctionalised plastid translocon regulates chloroplast ageing in *Arabidopsis*"

Bag et al., 2026

This file contains,

Supplementary tables 1 and 2

Supplementary figures 1-12

**Supplementary Tables:**

**Supplementary table 1: Primers used for this study**

The T-DNA and gene specific primers used for genotyping

| Gene | Primer sequences (5' to 3') |
|---|---|
| ***toc90-1*** | TAGCATCTGAATTTCATAACCAATCTCGATACAC LB3 (left border)<br>TTCCGTGACCCTCATCAAGAAC forward<br>GAGAAATCACTGTATCGCATGTCG reverse |
| ***toc90-2*** | TGGTTCACGTAGTGGGCCATCG LBa1 (left border)<br>GAGAAATCACTGTATCGCATGTCG reverse<br>TTCCGTGACCCTCATCAAGAAC forward |
| ***toc90-3*** | TGGTTCACGTAGTGGGCCATCG LBa1 (left border)<br>CAGCGCATGGATGTGGTTCAA forward<br>CTCCCGCCGTTTGCTGTTT reverse |
| ***toc34-2*** | TGGTTCACGTAGTGGGCCATCG LBa1 (left border)<br>TAATTTGATACGAGGTCAGCGAATCCGGC forward<br>TGATCTGGTTTGACTACACAATCGCACCC reverse |
| ***toc33*** | ATAACGCTGCGGACATCTAC LB/CGN3<br>GGTCTCTCGTTCGTGAATGG 127 Ex2 forward<br>CTGAGCGCCTATGATAAGAG 127 Ex7 reverse |
| ***toc132-2*** | CCCATTTGGACGTGAATGTAGACAC LB Gabi kat (left border)<br>GATGGGACTGAGTTTGTGGTTAGGTC forward<br>TCCTCCAACAATTGGTCTCTTGCTACCAG reverse |
| ***toc75-III-3*** | GCAAATCACAGGTGGACCAG dCAPS forward<br>TGCCCATGGAGGACTAGAAC dCAPS reverse |
| ***toc75-IV-1*** | GGTGCAGCAAAACCCACACTTTTACTTC dspm11<br>CTATCATCTTCTACTCCGAGTCTCCG forward<br>GCTGAATGAAACTTGTTAATGATAGT reverse |
| ***toc120-2*** | CAGTCATAGCCGAATAGCCTCTCCA FISH2<br>CTCCCAGAAAGCTGGATACTGTCAATGTG forward<br>AATTGGTTCGCAGGAGGGTC forward<br>CTTTCAGTCTCTCCCTTCTC reverse |
| ***toc159*** | GATGCAATCGATATCAGCCAATTTTAGAC LB102A<br>CAGTAGCAAAGCGGAAATGGACTCAAAG forward<br>GCACGTGGAGCTGCTGCAGGCTTCAAGGG reverse |

**(b)** Primers used in RT-PCR

| Primer name | Primer sequences (5’ to 3’) |
|---|---|
| ***eIF4E1*-F**<br>***eIF4E1*-R** | AAGATTTGAGAGGTTTCAAGCGGTGTAAG<br>AAACAATGGCGGTAGAAGACACTC |
| ***Toc90*-F**<br>***Toc90-R*** | CTCCCGCCGTTTGCTGTT<br>TTCCGTGACCCTCATCAAGAAC |
| ***Toc90*attB1**<br>***Toc90*attB2** | AAAAAGCAGGCTATGAAAGGCTTCAAAGACTG<br>AGAAAGCTGGGTTTAGGAAACGAGAAAATTCAC |
| ***Toc90* (H4C12)-F**<br>***Toc90* (H4C12)-R** | CATCTGGCATACCGGAGCTTGA<br>CTTCGAAACACGGCGGGACTT |
| **NTAPi-F**<br>**127 Ex2-F**<br>**Toc34 RT-R** | CATGCCATGGTGGTCGACAA<br>GGTCTCTCGTTCGTGAATGG<br>ACTTGCTAAACCGGAGTCTCG |

**Supplementary table 2:** Genetic analyses of *Arabidopsis* Toc90 double and triple mutants. The labelled column "phenotypic difference is stating which of these double and triple Toc90 mutants revealing any phenotypic difference from the control

| Genotype | Zygosity | Appropriate controls for phenotypic comparison | Phenotypic difference |
|---|---|---|---|
| *toc90-1 toc33* (*ppi1*) | hom/hom | *ppi1, toc90-1,* Col-0 | NO |
| *toc90-1 toc34-2* (*ppi3-2*) | hom/hom | *ppi3-2, toc90-1,* Col-0 | NO |
| *toc90-1 toc120-2* | hom/hom | *toc120-2, toc90-1,* Col-0 | NO |
| *toc90-1 toc132-2* | hom/hom | *toc132-2, toc90-1,* Col-0 | NO |
| *toc90-1 toc75-III-3* | hom/hom | *toc75-III-3, toc90-1,* Col-0 | NO |
| *toc90-1 toc75-IV-1* | hom/hom | *toc75-IV-1, toc90-1,* Col-0 | NO |
| *toc90-1 toc33 toc132-2* | hom/hom/hom | *toc33 toc132-2,* Col-0 | NO |
| *toc90-1 toc159 toc34-2* | hom/seg/hom | *toc159, toc34-2 toc159* | NO |
| *toc90-1 toc159 toc33* | hom/seg/hom | *toc159, toc33 toc159* | NO |
| *toc90-1 toc159* (*ppi2*) | hom/seg | *toc159* (albino plants) | YES |
| *toc90-1 toc132-2 toc120-2* | hom/hom/hom (bleached) | *toc132-2 toc120-2* (bleached) | YES |
| *toc90-1 toc132-2 toc120-2* | hom/hom/het (pale) | *toc132-2 toc120-2* (pale) | YES |
| *toc90-1 toc159 toc120-2* | hom/seg/hom | *toc90-1 toc159, toc159, toc159 toc120-2* | YES |
| *toc90-1 toc33 toc34-2* | hom/hom/het | *toc33 toc34-2* | YES |

Abbreviations in the table are defined as follows: hom, homozygous; het, heterozygous; seg, segregating; Col-0, Columbia

**Supplementary Figures:**

**Supplementary Figure 1.**

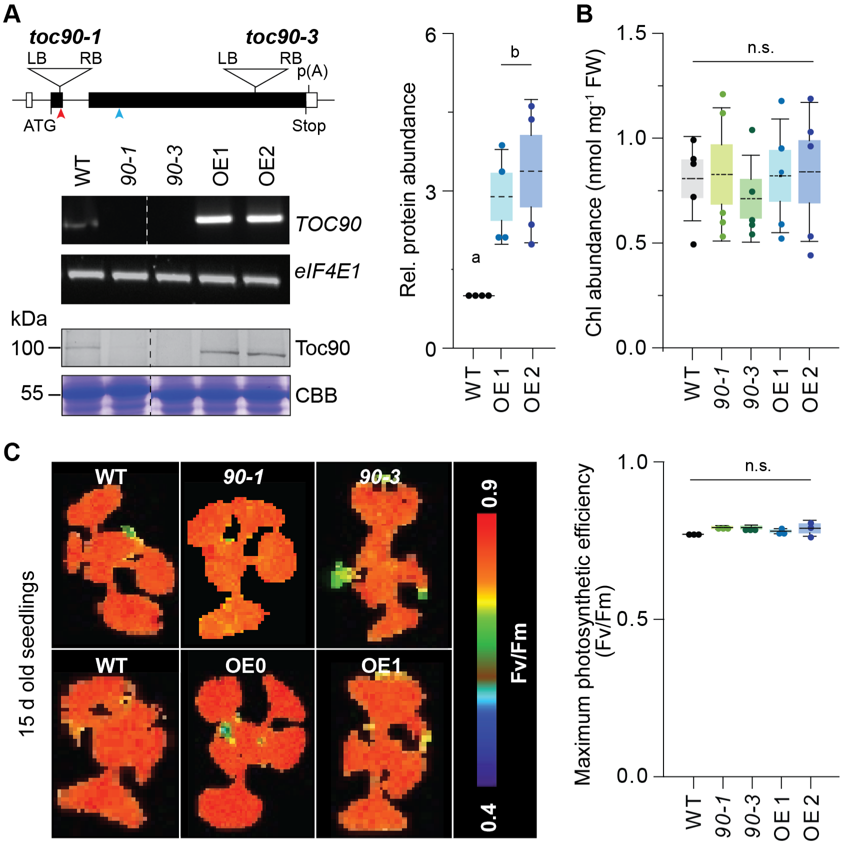


**Supplementary Fig 1. (A)** Schematic representation of the Arabidopsis thaliana Toc90 gene model. Represented are the sites of T-DNA insertion for the knockout lines *toc90-1* and *toc90-3*. The binding sites for the primers used for RT-PCR are represented by red and blue arrows. These primers were used to estimate the relative abundance of TOC90 mRNA levels in wild-type. Relative abundance of *TOC90* mRNA and Toc90 protein in wild-type (WT), knockout (*toc90-1*, *toc90-3*), and overexpression lines (OE1 and OE2). Eukaryotic Translation Initiation Factor 4E (*eIF4E*) used as control. PCR amplifications were performed using 23 cycles for *eif4e* and 32 cycles for *toc90*. Total protein (40 µg) was separated by SDS-PAGE and immunoblotted using a homemade polyclonal anti-TOC90 antibody (see Methods). A Coomassie Brilliant Blue (CBB) stained duplicate gel is shown as loading control. TOC90 protein levels were estimated by densitometry analysis of the western blots and normalised to WT levels (n=4). **(B)** Total chlorophyll abundance per gram fresh weight in the WT, KO and OE lines measured using DMF extraction from young (12-15 days) seedlings (n=5). **(C)** False colour image of maximum photosynthetic efficiency (Fv/Fm) and their absolute values in the WT, KO and OE lines (n=5). Statistical analysis in (A, B and C) was performed using one-way ANOVA followed by Fisher's least significant difference (LSD) test for pairwise comparisons among genotypes. Homogeneity of variance was assessed using Levene's test.

**Supplementary Figure 2.**

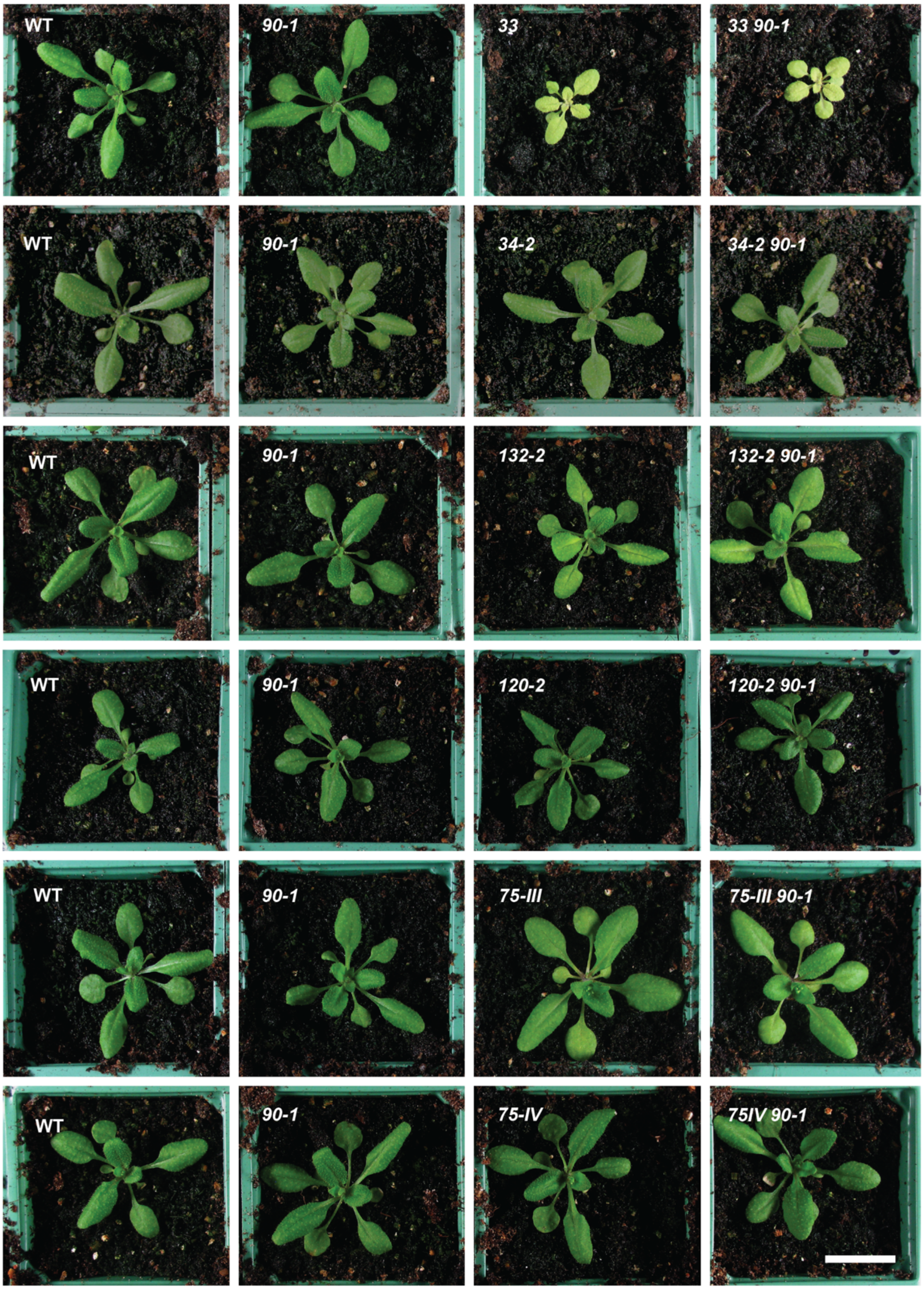


**Supplementary Fig 2.** Genetic interactions of the *toc90* knockout with other TOC component mutations. Representative phenotypes of *toc90-1* double mutants with *toc33, toc34-2*, *toc132-2*, *toc120-2*, *toc75-III-3*, and *toc75-IV-1* are shown alongside their respective parent single mutants and WT at 24 days after germination. Seeds were germinated in vitro on half strength of Murashige and Skoog (MS) medium and transferred to soil after 14 days of growth. Consistent with previous reports, *toc33* exhibits a pale, growth-retarded phenotype, while *toc34-2*, *toc132-2*, *toc120-2*, *toc75-III-3*, and *toc75-IV-1* display milder defects relative to WT. The *toc90-1* single mutant is indistinguishable from WT, and the *toc90-1* double mutants largely resemble their corresponding single mutant parents indicating that loss of TOC90 does not significantly enhance or suppress the phenotypes of other TOC component mutations. Scale bar = 1 cm.

**Supplementary Figure 3.**

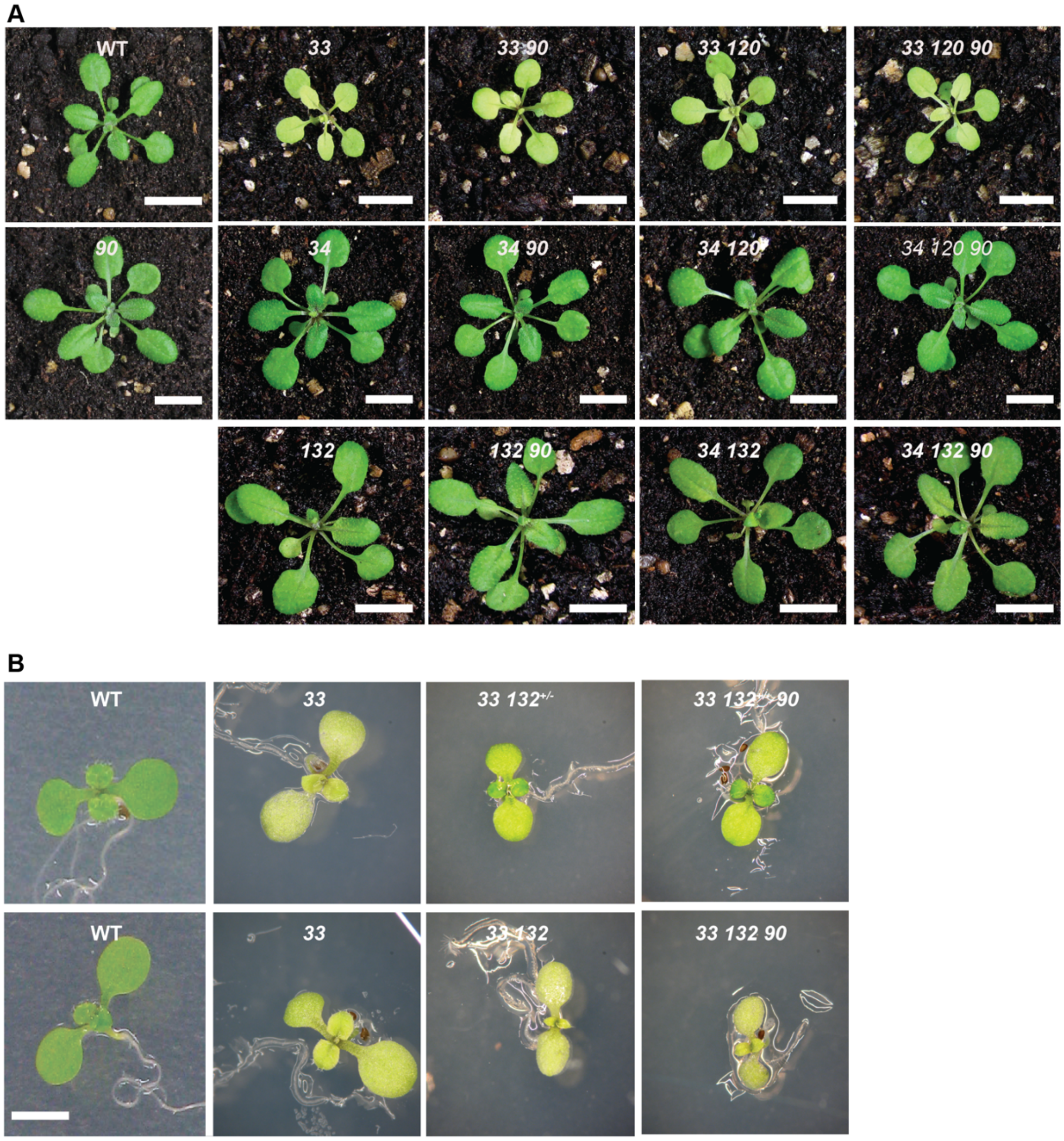


**Supplementary Figure 3. (A)** Phenotypic analysis of *toc90* triple homozygous mutants compared with the respective parental lines and WT. The *toc120 toc90 toc33*, *toc120 toc90 toc34*, and *toc132 toc90 toc34* triple mutants were germinated in vitro on half strength of Murashige and Skoog (MS) medium, transferred to soil after 14 days, and photographed at 24 days. In all cases, the triple mutants are indistinguishable from their respective single and double mutant parent lines and WT. Scale bars: 0.8 cm in the upper and lower panels and 1 cm in middle panel.
**(B)** Seedling-stage analysis of the *toc132 toc90 toc33* triple mutant compared with heterozygous and homozygous controls, grown in vitro for 8 days and imaged using a Zeiss dissecting microscope (10× magnification). No visible differences were observed at either early developmental stages or later growth. Scale bars: 2 mm.

**Supplementary Figure 4.**

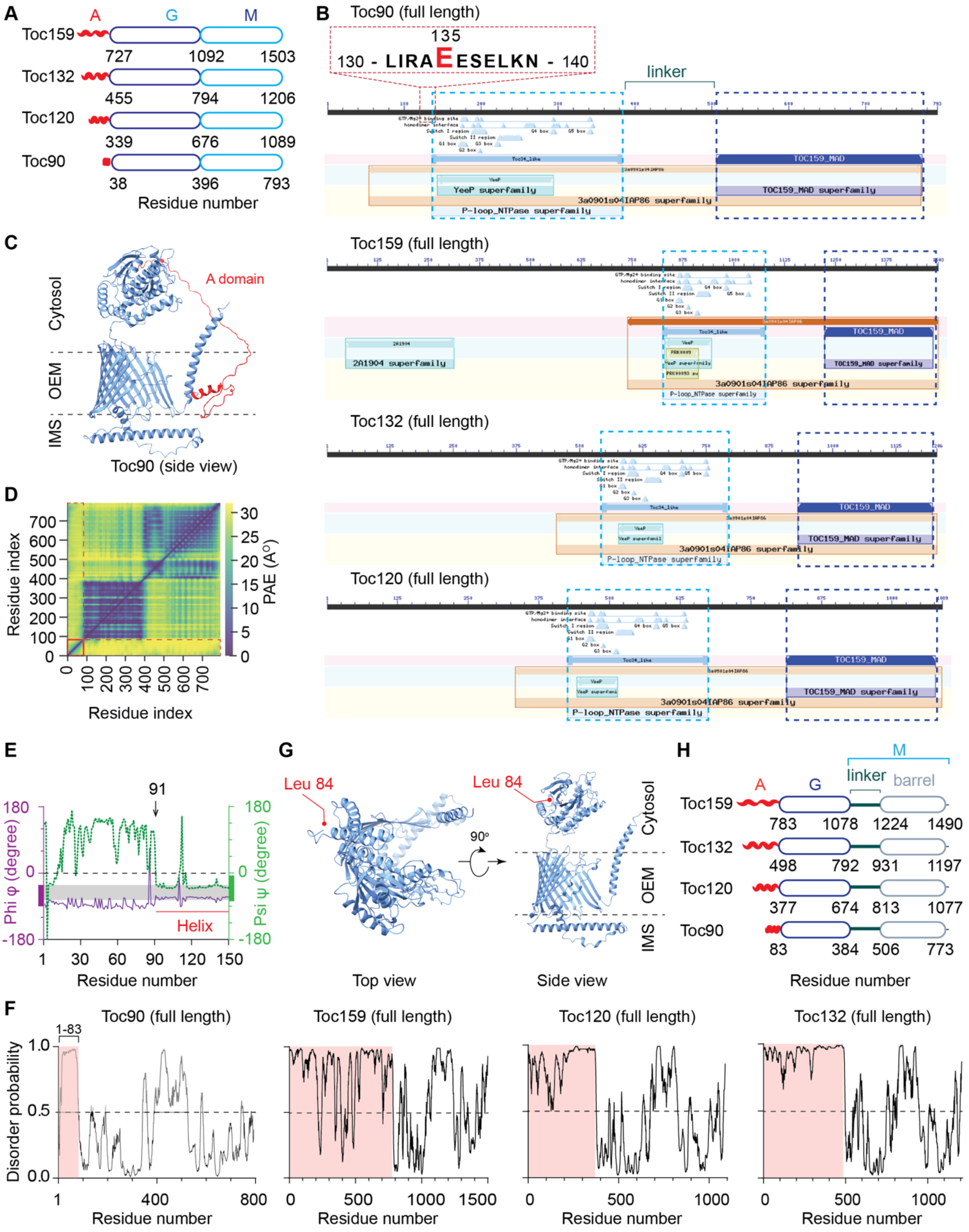


**Supplementary Fig 4. (A)** Current classification of A, G and M domains in the four GTPase receptors as per Ivanova et al., 2004. **(B)** Conserved domains prediction in the four *Arabidopsis* Toc homologues with NCBI CDD tool. All predictions were performed against CDD v3.21 - 62456 PSSMs database with default Expect Value (E-value) of 0.01. (The superfamily assignment follows the domain architecture described by Chen et al. (2000) and should be considered a curated representation rather than an exact reflection of domain boundaries predicted by the algorithm). The predicted start of the Toc34 like G-domain at E-135 shown in inset on top of Toc90 prediction. **(C)** AlphaFold-predicted structures of *Arabidopsis* Toc90 domains are colored according to NCBI CDD predictions: Acidic (red), GTPase and Membrane (steel blue). **(D)** Predicted alignment error in the AF3 predicted Toc90 structure. **(E)** Phi (φ) and Psi (Ψ) dihedral angle of first 150 amino acid residues of mature Toc90 protein. **(F)** Disorder probability score of the Acidic domains of all four *Arabidopsis thaliana* Toc159 family proteins. **(G)** Acidic domain removed Toc90 structure top and side view. First residue (Leucine) colored in red. **(H)** Classification of A, G and M domain of all the Toc159 family proteins as deduced from structural and bioinformatic analysis.

**Supplementary Figure 5.**

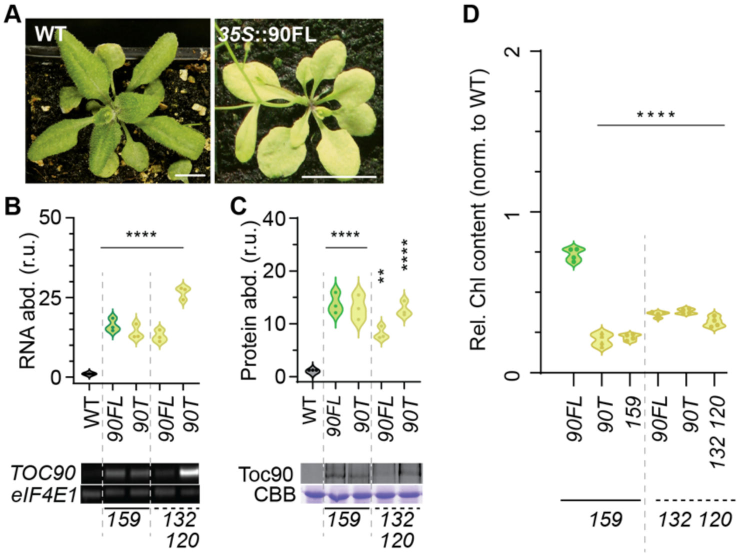


**Supplementary Fig 5. (A)** Representative soil-grown WT (Scale bar 0.5 cm) and toc159 knockout plants expressing *35S*::90FL (1 cm). **(B)** Relative *TOC90* transcript abundance in wild type and independent transgenic *35S*::90FL lines generated in the two mutant backgrounds. Total RNA was extracted from 14-day-old in vitro-grown seedlings, and transcript accumulation was assessed by semi-quantitative RT-PCR using Toc90-specific primers (Supplementary table 1b) designed to detect transgene overexpression. *eIF4E1* was used as an internal control. PCR amplification was terminated after 26 cycles for Toc90 and 22 cycles for *eIF4E1*. Representative agarose gel images are shown below the quantification, and band intensities were normalised to *eIF4E1*. **(C)** Relative Toc90 protein abundance in the same genotypes. Protein levels were quantified from immunoblots and normalised to total protein loading (30 µg per lane), shown by Coomassie Brilliant Blue (CBB) staining. **(D)** Relative chlorophyll content normalised to WT in *toc159* knockout and *toc132 toc120* knockout backgrounds expressing Toc90 constructs or corresponding control lines. Dots represent individual biological replicates, and violin plots show the distribution of the data. Asterisks indicate statistically significant differences relative to the WT or indicated control (*35S*::90FL in D) comparison: P < 0.01, ****P < 0.0001.

**Supplementary Figure 6.**

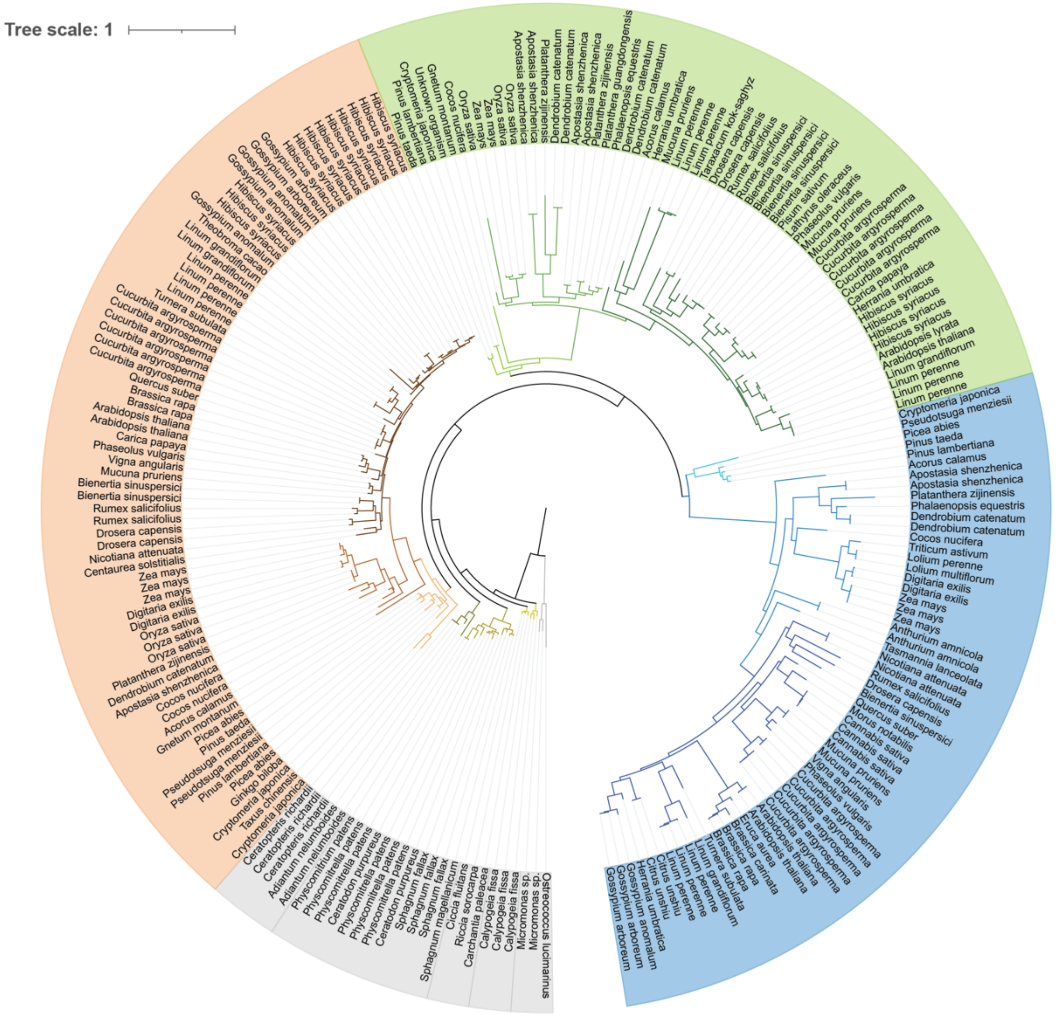


**Supplementary Fig 6.** Fully annotated phylogenetic tree with all 262 full length TOC sequences across the 67 species. Shaded regions around species names indicate TOC classification: grey - unclassified, salmon - Toc120, green - TOC159 and blue - TOC90. Tree branch colour corresponds to the branch colours in the main figure 3.

**Supplementary Figure 7.**

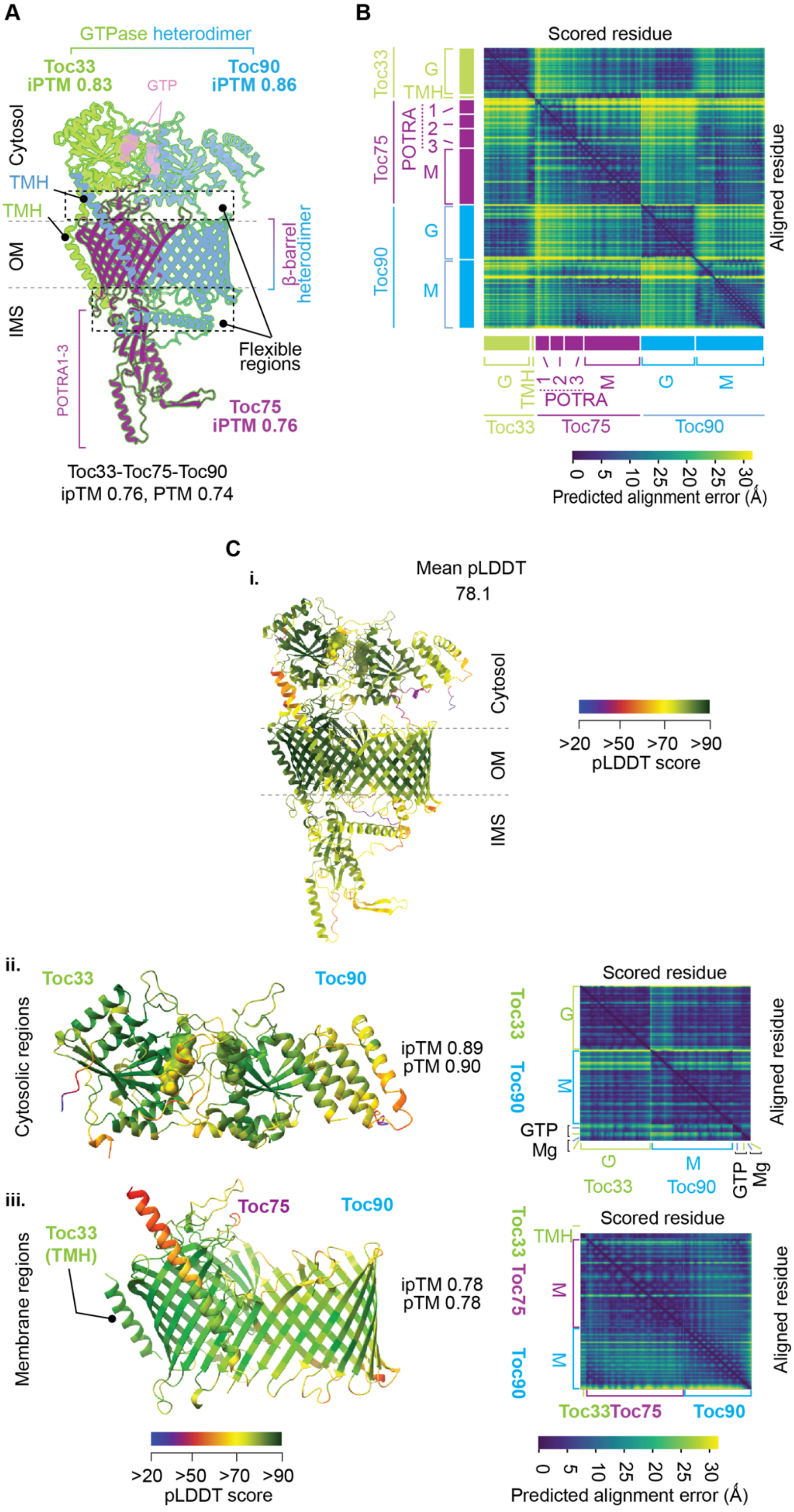

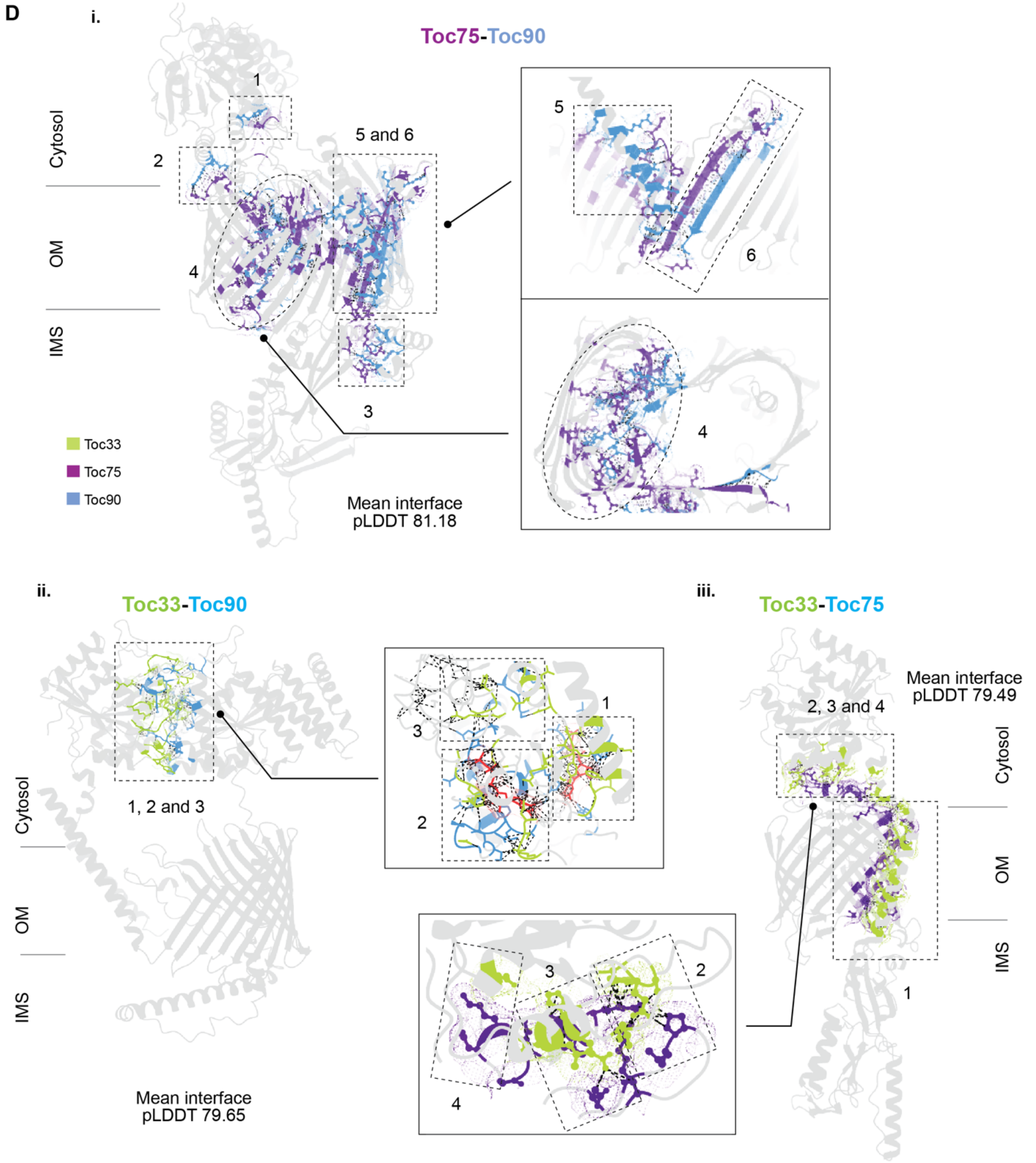


**Supplementary Fig. 7. (A)** AF3-predicted model of the Toc90 complex. Domains are annotated according to refs. 18, 19 and 40. Chain-wise average ipTM scores are shown for the individual components. **(B)** Predicted alignment error (PAE) of the modelled complex, shown as a heat map in which residue positions are plotted against aligned residue positions. The corresponding domains and subdomains of each protein are annotated along both axes. **(C)** Residue-wise colour coding according to pLDDT score for (i) the complete complex, (ii) the cytoplasmic module and (iii) the membrane module, together with their corresponding average pLDDT scores. The overall ipTM and pTM scores are also indicated. Module-specific pLDDT values are provided below as additional quality-control measures. Collectively, these scores indicate high structural confidence for the predicted model. **(D)** Pairwise average interface and contact-site pLDDT scores for the predicted interactions within the complex, highlighting regions of high structural confidence and potential interaction surfaces of further interest. **(i)** Toc75-Toc90 interface. Six predicted contact sites (1-6) are identified between Toc75 and Toc90. Sites 1 and 2 involve interactions between helices of the Toc90 G-domain and flexible loops connecting the β-strands forming the Toc75 β-barrel on the cytoplasmic side. Site 4 shows an interaction between Toc90 and the interior of the Toc75 β-barrel, involving loops associated with the Toc90 membrane region and the inner surface of Toc75 (See further in **Supplementary Fig. 8E**). Site 5 involves interaction of the Toc90 transmembrane helix (TMH) with the outer surface of the Toc75 β-barrel. Site 6 involves an interaction between Toc90 β-strand 14 and Toc75 β-strand 17, the latter forming part of the flap proposed to regulate pore size and opening of the Toc75-containing β-barrel assembly. **(ii)** Toc33-Toc90 interface. Three predicted contact sites (1-3) are identified between the cytoplasmic G-domains of Toc33 and Toc90. Sites 1 and 2 are shown in greater detail in Fig. 4B and C in relation to GTP binding and the predicted G-domain interface. Site 3 involves interactions between loop regions of the two G-domains located largely outside the catalytic nucleotide-binding pockets. **(iii)** Toc33-Toc75 interface. Four predicted contact sites (1-4) are identified between Toc33 and Toc75. Site 1 involves interaction between the Toc33 TMH and the Toc75 β-barrel. Sites 2-4 involve interactions between helices of the Toc33 G-domain and flexible loops connecting the β-strands forming the Toc75 β-barrel on the cytoplasmic side.

**Supplementary Figure 8.**

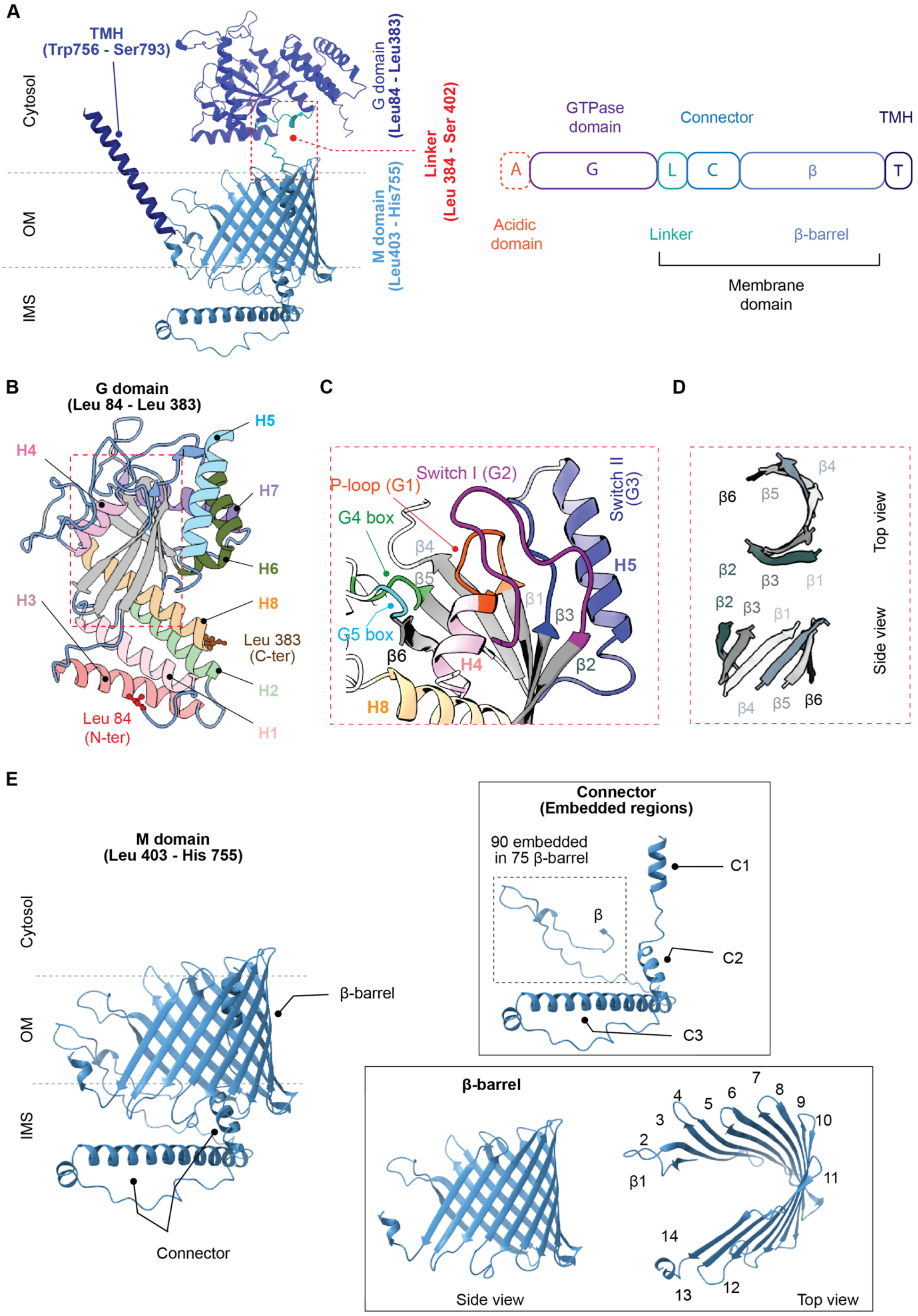


**Supplementary Fig 8. (A)** Schematic and overall structural organisation of Toc90, highlighting the major predicted domain features including A domain, G domain, Linker, Connector and M domain. Note A domain was not included in the structural model. **(B)** Structural features of the Toc90 G-domain, with emphasis on the arrangement of α-helices surrounding the catalytic core. **(C)** β-sheet architecture within the Toc90 G-domain, showing the central β-sheet framework characteristic of GTPase domains. **(D)** Positioning of the conserved GTPase catalytic motifs, including the G1-G5 boxes, within the Toc90 G-domain. These motifs define the nucleotide-binding and catalytic region of the protein. **(E)** Structural features of the Toc90 M-domain components, including the embedded connector region and the β-barrel-like domain. These elements form the C-terminal portion of Toc90 and may contribute to the spatial organisation and membrane-associated features of the protein.

**Supplementary Figure 9.**

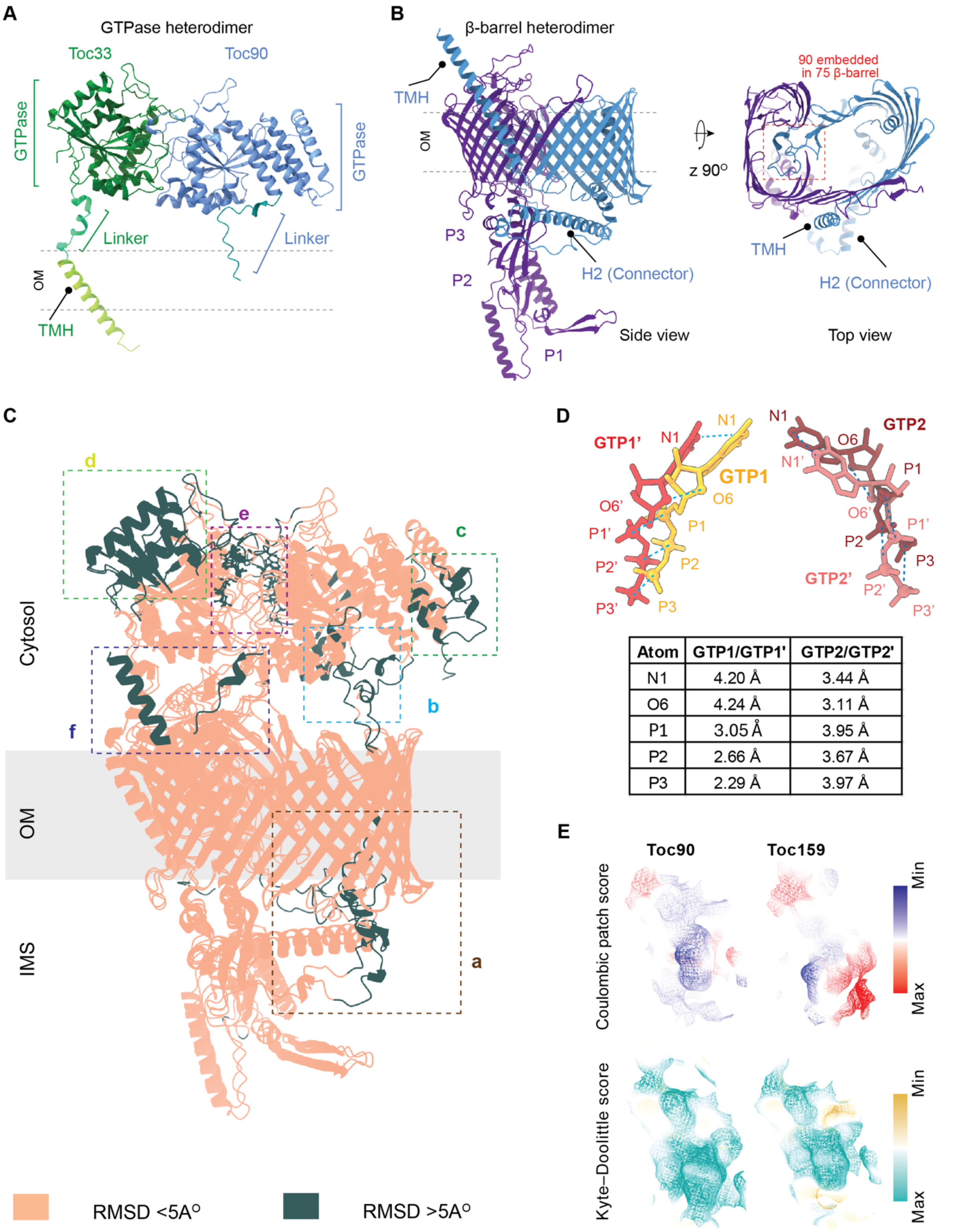


| Atom | GTP1/GTP1' | GTP2/GTP2' |
|---|---|---|
| N1 | 4.20 Å | 3.44 Å |
| O6 | 4.24 Å | 3.11 Å |
| P1 | 3.05 Å | 3.95 Å |
| P2 | 2.66 Å | 3.67 Å |
| P3 | 2.29 Å | 3.97 Å |

**Supplementary Fig. 9. (A)** GTPase heterodimer formed by Toc33 and the Toc90 G-domain, together with the associated linker and transmembrane helix (TMH) regions. The portion of the Toc90 linker that remains embedded within the Toc75 β-barrel is indicated by the red box. **(B)** β-barrel heterodimer formed by Toc75 and Toc90, showing the associated POTRA and TMH regions in side and top views. **(C)** Superposition of the Toc33-Toc75-Toc90 and Toc33-Toc75-Toc159 complexes. Regions in which the Cα–Cα distance between corresponding residues is <5

Å are shown in salmon pink, whereas regions with distances >5 Å are shown in dark green. Six major regions displaying deviations >5 Å are labelled a–f and highlighted by coloured boxes for clarity. **(D)** Comparison of the predicted orientations of the two GTP molecules in the Toc90 complex (GTP1 and GTP2) and the Toc159 complex (GTP1′ and GTP2′). The table below shows the distances between corresponding atoms of the aligned GTP molecules in the two complexes. **(E)** Electrostatic surface properties, represented by Coulombic potential, and hydrophobicity, represented by the Kyte-Doolittle scale, are shown for the nucleotide-binding pockets of Toc90 and Toc159.

**Supplementary Figure 10.**

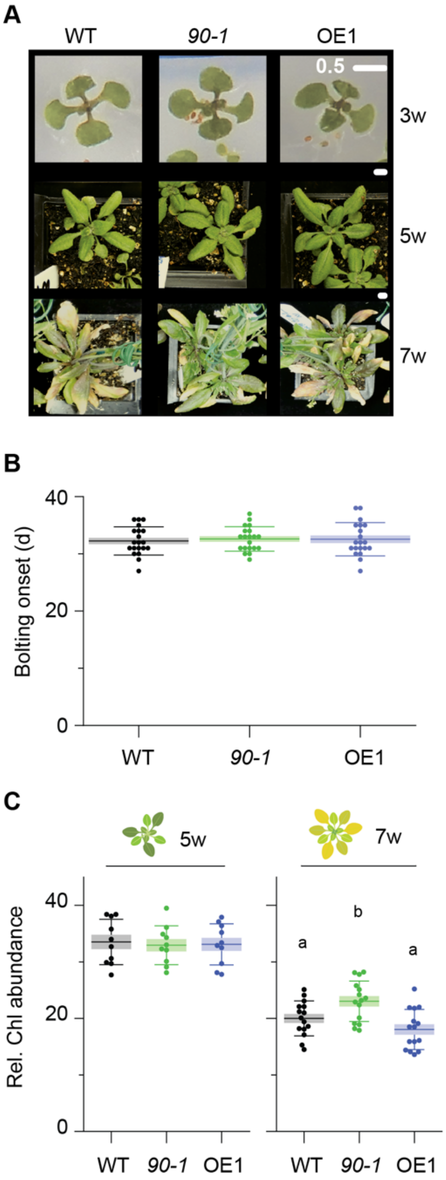


**Supplementary Fig 10. (A)** Pilot phenotypic characterisation of WT, *toc90-1*, and OE1 plants at 3, 5, and 7 weeks after emergence. This pilot experiment was performed once, with more than 30 plants per genotype assessed in parallel for the different phenotypic measurements. The experiment was intended as an exploratory screen to identify whether Toc90 loss or overexpression caused any obvious developmental phenotype, rather than as a definitive replicated phenotyping assay. Since clear developmental differences were observed at 7 weeks old plants, the experiment was not repeated as a full-scale phenotypic study, but more systematic experimental approaches were taken as in Fig 5B-F. Scale bar = 0.5 cm. **(B)** Bolting time in WT, *toc90-1*, and OE1 plants (n=12). Bolting time is ideally quantified using rosette leaf number in addition to chronological age. However, because no developmental differences were observed during the first 3 weeks, any potential shift in the onset of bolting was likely to be small, possibly only a few days. Such a subtle difference may not be sufficient to alter rosette leaf number, but it could become apparent when bolting is assessed in a larger population and at higher temporal resolution. Thus, the absence of an obvious bolting phenotype does not exclude a subtle developmental role for Toc90 but suggests that Toc90 is not essential for vegetative development or for the major transition to flowering under the conditions tested. **(C)** Chlorophyll abundance in WT, *toc90-1*, and OE1 plants at 5 and 7 weeks after emergence (n=10) expressed in terms of SPAD values. No difference in chlorophyll abundance was observed at 3 weeks after emergence, as shown in Supplementary Fig. 1B. Statistical analysis in (A and B) was performed using one-way ANOVA followed by Fisher's least significant difference (LSD) test for pairwise comparisons among genotypes. Homogeneity of variance was assessed using Levene's test.

**Supplementary Figure 11.**

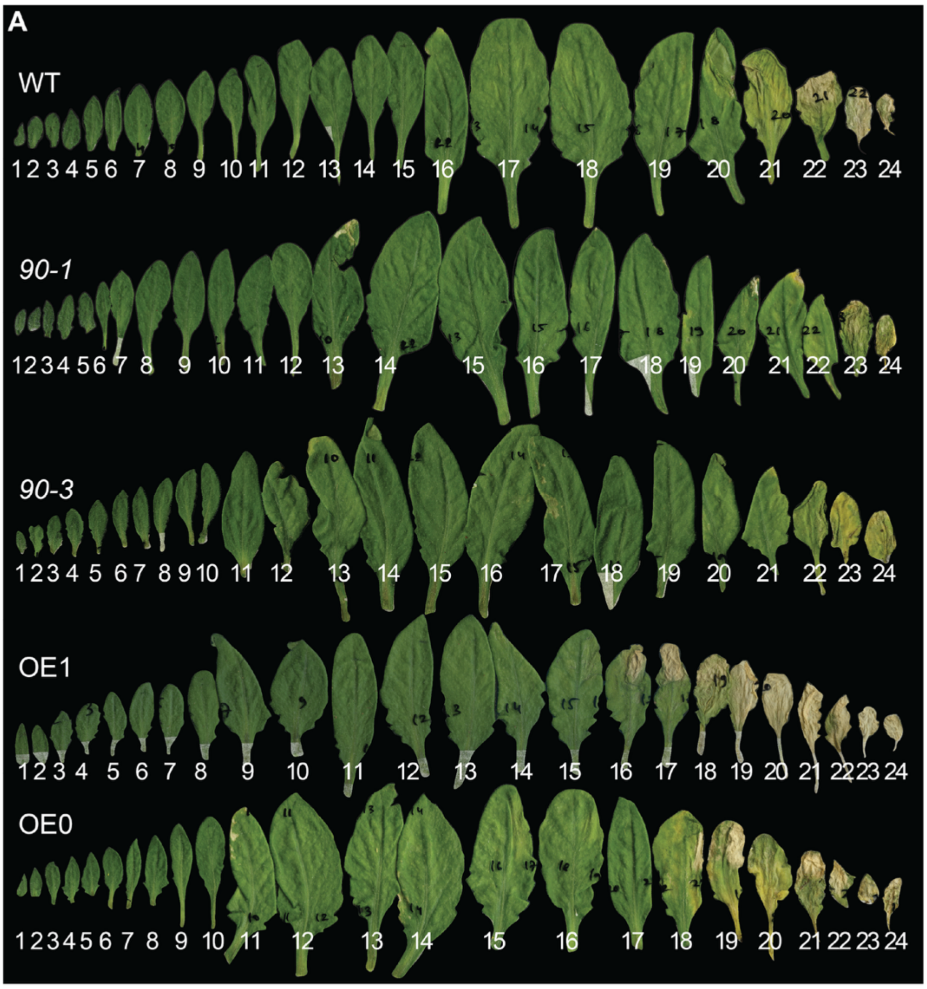


**Supplementary Fig 11.** Fully opened rosette image from Fig 5B. Note that leaf numbers here are done in a reverse order.

**Supplementary Figure 12.**

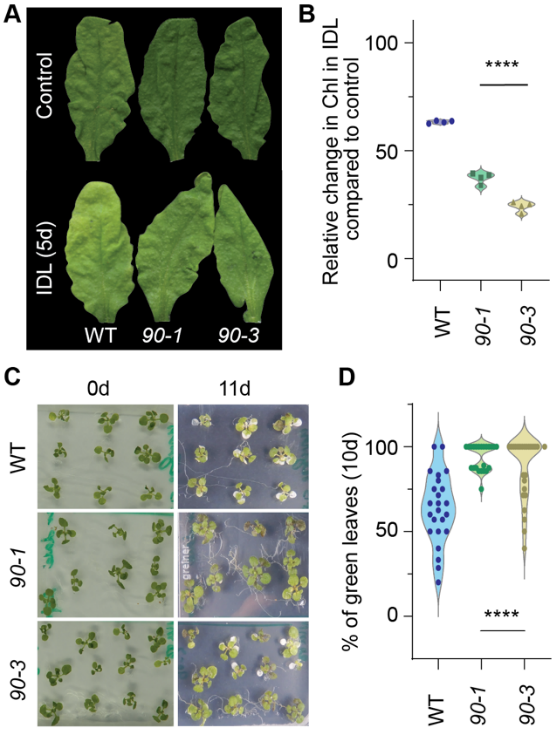


**Supplementary Fig 12. (A)** Dark-induced senescence in WT and *toc90* lines after 5 days of darkness, compared with non-dark-treated leaves (IDL - individually darkened leaves). **(B)** Relative change in chlorophyll abundance after dark treatment, normalised (100%) to the corresponding non-dark-treated leaves in WT and *toc90* lines. **(C)** Paraquat treatment assay in WT and *toc90* seedlings. Seedlings were germinated on medium without paraquat, transferred to plates supplemented with 10 µM paraquat, and imaged after 11 days under long-day growth conditions. **(D)** Quantitative analysis of paraquat sensitivity, expressed as the percentage of green leaves without visible white spots caused by paraquat stress (n=4 independent experiments).
